# *Plagl1* and *Lrrc58* control mammalian body size by triggering target-directed microRNA degradation of miR-322 and miR-503

**DOI:** 10.1101/2025.06.30.662380

**Authors:** Collette A. LaVigne, Jaeil Han, He Zhang, Sihoon Cho, Minseon Kim, Komal Sethia, Bret M. Evers, Asha Acharya, Tsung-Cheng Chang, Joshua T. Mendell

**Affiliations:** Department of Molecular Biology, University of Texas Southwestern Medical Center, Dallas, TX 75390, USA; Department of Microbiology and Molecular Biology, Chungnam National University, Daejeon, Republic of Korea; Department of Health Data Science and Biostatistics, Peter O’Donnell Jr. School of Public Health, University of Texas Southwestern Medical Center, Dallas, TX 75390, USA; Department of Microbiology, University of Texas Southwestern Medical Center, Dallas, TX 75390, USA; Department of Pathology, University of Texas Southwestern Medical Center, Dallas, TX 75390, USA; Department of Ophthalmology, University of Texas Southwestern Medical Center, Dallas, TX 75390, USA; Harold C. Simmons Comprehensive Cancer Center, University of Texas Southwestern Medical Center, Dallas, TX 75390, USA; Hamon Center for Regenerative Science and Medicine, University of Texas Southwestern Medical Center, Dallas, TX 75390, USA; Howard Hughes Medical Institute, University of Texas Southwestern Medical Center, Dallas, TX 75390, USA

## Abstract

Precise control of microRNA (miRNA) expression is critical during development. An important mechanism of miRNA regulation is target-directed microRNA degradation (TDMD), a pathway in which the binding of miRNAs to specialized trigger RNAs induces ubiquitylation and decay of associated Argonaute (AGO) proteins by the ZSWIM8 ubiquitin ligase. Concomitant release of miRNAs results in their rapid turnover. ZSWIM8-deficient mice exhibit reduced body size, cardiopulmonary and neurodevelopmental defects, and perinatal lethality. Despite widespread dysregulation of miRNAs in these animals, the vast majority of presumptive trigger RNAs that induce decay of ZSWIM8-regulated miRNAs remain undefined. Here, using AGO crosslinking and sequencing of hybrids (AGO-CLASH), a high-throughput method for identifying miRNA binding sites, we report the identification of *Plagl1* as a TDMD trigger for miR-322-5p, and *Lrrc58* and *Malat1* as TDMD triggers for miR-503-5p in mouse embryonic fibroblasts (MEFs). In mice, deletion of the miR-322-5p and miR-503-5p trigger sites in the *Plagl1* and *Lrrc58* 3′ UTRs, respectively, abrogated TDMD of these miRNAs and resulted in miR-322/503-dependent embryonic growth restriction, recapitulating a key feature of the *Zswim8^—/—^* phenotype. Thus, *Plagl1* and *Lrrc58* act as triggers for degradation of miR-322-5p and miR-503-5p, revealing a noncoding function for these mRNAs as regulators of mammalian body size.

## INTRODUCTION

microRNAs (miRNAs) are small noncoding RNAs that function as important post-transcriptional regulators of gene expression across eukaryotes (Bartel 2018). miRNAs are produced by a highly regulated biogenesis pathway that processes long primary miRNA transcripts (pri-miRNAs) into short ∼22 nucleotide double-stranded RNA duplexes (Kim et al. 2025). One strand of the duplex, representing the mature miRNA, is loaded into an Argonaute protein (AGO), while the other strand, referred to as the passenger strand or miRNA*, is degraded. miRNA loaded-AGO proteins surveil mRNA and noncoding RNA transcripts for sites with complementarity to nucleotides 2-7 of the miRNA, termed the seed sequence. Base pairing between the miRNA and target RNA enables AGO to recruit deadenylation and decapping factors, ultimately leading to repression of translation and/or accelerated degradation of the target (Jonas and Izaurralde 2015; Shang et al. 2023). The majority of mammalian genes have at least one conserved miRNA binding site (Friedman et al. 2009), allowing miRNAs to broadly sculpt gene expression during development and disease (Mendell and Olson 2012; Alberti and Cochella 2017; DeVeale et al. 2021). Consequently, elaborate mechanisms that control miRNA transcription, processing, and degradation have evolved in order to precisely control the timing and magnitude of gene silencing by this pathway (Winter et al. 2009; Gebert and MacRae 2019; Hiers et al. 2024).

The loading of a miRNA into an AGO protein shields it from degradation, usually conferring a long half-life that enables many cycles of target RNA repression (van Rooij et al. 2007; Gatfield et al. 2009). However, analyses of miRNA decay rates revealed a set of exceptional miRNAs that exhibit accelerated turnover (Hwang et al. 2007; Bail et al. 2010; Krol et al. 2010; Gantier et al. 2011; Rissland et al. 2011; Guo et al. 2015; Marzi et al. 2016; Kingston and Bartel 2019; Reichholf et al. 2019). It is now appreciated that a mechanism called target-directed miRNA degradation (TDMD) accounts for the enhanced decay of many of these short-lived miRNAs.

TDMD is activated upon base pairing of a miRNA to a specialized mRNA or noncoding RNA, referred to as a trigger RNA (Ameres et al. 2010; Cazalla et al. 2010). Our understanding of the base-pairing architecture, and potentially other features, that differentiate a canonical miRNA target from a TDMD trigger is still evolving. The known trigger RNAs in *Drosophila* and mammals engage in extensive base pairing with both the seed region and 3′ end of the miRNA (Bitetti et al. 2018; Ghini et al. 2018; Kleaveland et al. 2018; Li et al. 2021; Kingston et al. 2022; Sheng et al. 2023), while in *C. elegans*, evidence suggests that base-pairing of the miRNA seed is sufficient to initiate TDMD in a subset of cases (Donnelly et al. 2022; Stubna et al. 2024).

TDMD requires a cullin-RING ubiquitin ligase complex containing the adapter protein ZSWIM8 (Han et al. 2020; Shi et al. 2020). Upon binding of a miRNA to a trigger, the ZSWIM8 complex is believed to ubiquitylate the associated AGO protein, leading to proteasome-mediated degradation and concomitant release and decay of the miRNA. Extensive base pairing between the miRNA and trigger, involving both the seed and miRNA 3′ end, results in a unique AGO conformation (Sheu-Gruttadauria et al. 2019) that is thought to be specifically recognized by ZSWIM8. How ZSWIM8 orthologs are activated in cases where accelerated miRNA degradation depends only on the seed sequence remains a mystery (Donnelly et al. 2022).

Identification of the protein machinery that mediates TDMD, and the deep conservation of ZSWIM8, has allowed investigation of the physiologic role and scope of miRNA regulation by this pathway in diverse metazoan species. In *C. elegans*, loss of the ZSWIM8 ortholog EBAX-1 causes defects in locomotion, egg-laying, male mating behaviors, and axon guidance (Wang et al. 2013). Small RNA sequencing documented more than 20 miRNAs that are stabilized in EBAX-1-deficient worms (Donnelly et al. 2022; Stubna et al. 2024). Dora, the *Drosophila* ortholog of ZSWIM8, is essential, with most *dora* knockout embryos failing to hatch into L1 larvae (Kingston et al. 2022; Molina-Pelayo et al. 2022). More than 20 Dora-regulated miRNAs have been identified in S2 cells and embryos (Shi et al. 2020; Kingston et al. 2022). Importantly, embryonic lethality in flies is partially rescued by reducing miR-3 levels, thereby directly implicating miRNA dysregulation in this aspect of the Dora-deficiency phenotype (Kingston et al. 2022). In mice, loss of ZSWIM8 results in reduced body size, cardiopulmonary and neurodevelopmental defects, and perinatal lethality (Wang et al. 2022; Jones et al. 2023; Shi et al. 2023). Small RNA sequencing of E18.5 embryonic mouse *Zswim8^—/—^* tissues revealed more than 50 upregulated miRNAs, suggesting widespread regulation of miRNAs by TDMD during mammalian development (Jones et al. 2023; Shi et al. 2023). Importantly, reduced dosage of miR-322-5p and miR-503-5p, two miRNAs that are co-transcribed as part of a miRNA cluster and share a six-nucleotide core seed sequence, rescued embryonic growth deficiency in *Zswim8^—/—^*embryos, confirming that this phenotype is caused by aberrant overexpression of these miRNAs (Jones et al. 2023).

While there now exists an extensive catalog of miRNAs that are regulated by ZSWIM8 orthologs and are therefore inferred to be TDMD substrates, the presumptive trigger RNAs that are responsible for initiating degradation of these miRNAs have proven more difficult to identify. In *Drosophila*, bioinformatic analyses seeking conserved miRNA binding sites with extensive 3′ complementarity led to the identification of six TDMD triggers that function in S2 cells or fly embryos (Kingston et al. 2022). Recently, crosslinking and sequencing of hybrids to identify AGO-associated miRNAs and their bound target RNAs (AGO-CLASH) (Helwak and Tollervey 2014) has emerged as a complementary tool to identify TDMD triggers. For example, AGO1-CLASH in *Drosophila* S2 cells led to the identification of the same TDMD triggers found by Kingston et al. (Sheng et al. 2023), supporting the use of this approach for identifying new triggers in additional settings.

The vast majority of presumptive trigger RNAs in mammals remain to be defined. Despite the existence of more than 50 known miRNAs that are destabilized by ZSWIM8 in mouse tissues (Jones et al. 2023; Shi et al. 2023), and bioinformatic predictions of hundreds of potential mammalian trigger sites (Simeone et al. 2022), only four endogenous trigger sites have been validated thus far (Bitetti et al. 2018; Ghini et al. 2018; Kleaveland et al. 2018; Li et al. 2021). The most recently discovered mammalian TDMD trigger, *BCL2L11*, was discovered by analyzing AGO-CLASH data generated using mammalian cell lines and tissues (Li et al. 2021). Given recent advances in the AGO-CLASH methodology that enable the enhanced recovery of chimeras containing specific miRNAs of interest (Manakov et al. 2022), we reasoned that additional AGO-CLASH experiments focused on miRNAs that are strongly regulated by ZSWIM8 in mouse tissues might uncover additional TDMD triggers. Here we report the successful use of this approach to identify trigger sites that are required for TDMD of miR-322-5p and miR-503-5p in mouse embryonic fibroblasts (MEFs) (*Plagl1*:miR-322-5p, *Lrrc58*:miR-503-5p, and *Malat1*:miR-503-5p). In mice, deletion of the miR-322-5p and miR-503-5p trigger sites in *Plagl1* and *Lrrc58* impaired TDMD of these miRNAs *in vivo* and resulted in embryonic growth deficiency, partially phenocopying a key characteristic of *Zswim8^—/—^* mice. This study therefore provides a valuable dataset that will facilitate identification of additional mammalian TDMD triggers and establishes the existence of a *Plagl1*/*Lrrc58*-mediated TDMD pathway that plays a major role in regulating mammalian body size.

## RESULTS

### AGO-CLASH identifies potential TDMD trigger RNAs

In order to identify new TDMD triggers, we focused on a set of 10 miRNAs that are strongly regulated by ZSWIM8 in both mouse tissues and contact-inhibited MEFs (Han et al. 2020; Shi et al. 2020; Jones et al. 2023; Shi et al. 2023) (Supplemental Fig. S1A). This set included miR-7a-5p as a positive control since its trigger RNA, *Cyrano* (also known as *Oip5-AS1*), has been identified (Kleaveland et al. 2018). We also included miR-322-5p and miR-503-5p because of their documented roles in regulating mammalian body size (Jones et al. 2023). A limitation of the traditional AGO-CLASH method is that it produces chimeric reads that are heavily biased towards highly abundant miRNAs and their targets. We therefore applied a recently developed AGO-CLASH protocol, also known as chimeric eCLIP, that incorporates an enrichment step using biotinylated probes to increase the representation of chimeras containing specific miRNAs of interest (Manakov et al. 2022). Two probes were designed to enrich for five miRNAs each (Fig. 1A and Supplemental Fig. S1A) and libraries were prepared using brain and lung tissue from wild type and *Zswim8^—/—^*mice (Jones et al. 2023). As expected, a substantial increase in chimeric reads corresponding to the enriched miRNAs was observed in both brain and lung (Fig. 1A). A large majority of chimeric reads comprising a miRNA and a target RNA with a corresponding seed match mapped to 3′ UTRs and coding sequences (Fig. 1B and Supplemental Fig. 1B), consistent with the known locations of *bona fide* miRNA binding sites (Bartel 2018).

**Figure 1.**
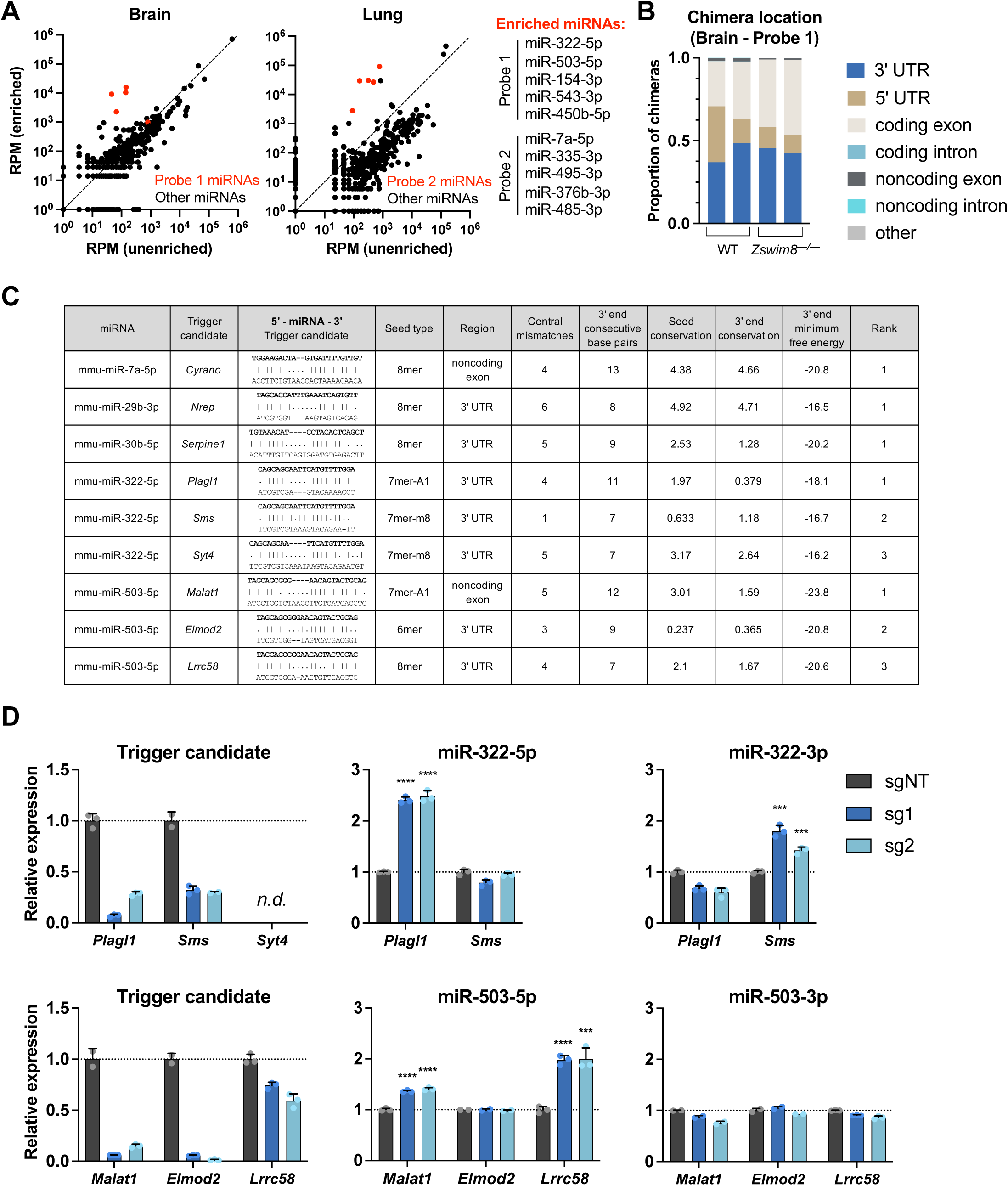
AGO-CLASH identifies potential TDMD trigger RNAs. (*A*) Plots showing reads per million (RPM) of the sum of all chimeras detected for each miRNA in enriched versus unenriched AGO-CLASH brain and lung samples. Enriched miRNAs shown in red. (*B*) Proportion of chimeras mapped to each location in replicate WT and *Zswim8^—/—^* brain AGO-CLASH samples enriched with Probe 1. Only chimeras in which the predicted base pairing to the miRNA seed sequence was a 6mer, 7mer-A1, 7mer-m8, or 8mer were included in the analysis. (*C*) Table of filtered and ranked candidate TDMD trigger sites identified by AGO-CLASH. Shown are the top ranked trigger for miR-7a-5p, miR-29b-3p, and miR-30b-5p, and the top three candidate triggers for miR-322-5p and miR-503-5p. (*D*) qRT-PCR analysis of candidate trigger sites in MEFs. Immortalized MEFs expressing dCas9-KRAB were infected with lentivirus encoding a non-targeting guide (sgNT) or two independent guides targeting candidate trigger RNAs (sg1, sg2). Shown is the candidate trigger expression normalized to *Actb* (left), mature miRNA abundance normalized to miR-16-5p (middle), and the passenger strand levels normalized to miR-16-5p (right). Values were normalized to expression level in sgNT for each transcript. n=3 technical replicates per sgRNA with individual data points plotted (mean ± SD shown). *P* values were calculated by one-tailed student’s t-test comparing sg1 or sg2 to sgNT. \*\*\**P*<0.001; \*\*\*\**P*<0.0001; n.d., not reliably detected.

Chimeric reads in these datasets are expected to predominantly correspond to canonical miRNA target sites, with only a small fraction representing TDMD trigger sites. Therefore, as an initial approach to nominate sites that are most likely to function as TDMD triggers, we developed a set of highly stringent filtering criteria based on the features exemplified by the existing collection of validated *Drosophila*, mammalian, and viral trigger RNAs (Buhagiar and Kleaveland 2024). First, we mandated the presence of a seed match that was classified as a 6mer, 7mer-A1, 7mer-m8, or 8mer (Lewis et al. 2005; Grimson et al. 2007), excluding any potential triggers with mismatches within the seed binding sequence. Second, we selected only those candidate sites that fell within noncoding RNAs or within noncoding regions of mRNAs (5′ or 3′ UTRs), based upon the observation that translating ribosomes diminish TDMD when the trigger site is located within an open reading frame (Li et al. 2025). Third, we excluded any potential trigger sites with large central bulges, defined as more than seven consecutive mismatches between the putative trigger and the 3′ sequence of the miRNA after the seed.

Fourth, we required the presence of at least six consecutive base pairs between the trigger and the 3′ region of the miRNA (defined as the final 13 nucleotides of the miRNA). Although mutagenesis experiments suggest that this latter feature is not absolutely required for TDMD (Han et al. 2020), all 13 validated TDMD triggers exhibit this attribute (Buhagiar and Kleaveland 2024). Fifth, we selected candidates that exhibit detectable evolutionary conservation of the seed binding region and the nucleotides predicted to base-pair with the 3′ region of the miRNA, as defined by a positive PhyloP (60 vertebrate) score (Pollard et al. 2010; Perez et al. 2025).

Last, the minimum free binding energy of the predicted duplex between the potential trigger and 3′ region of the miRNA (3′ MFE) was calculated and candidates were ranked based upon this metric. Importantly, this pipeline ranked the known TDMD triggers *Cyrano*, *Nrep*, and *Serpine1* as the top candidates for their respective miRNAs: miR-7a-5p, miR-29b-3p, and miR-30b-5p (Fig.1C). Thus, although these criteria are not expected to capture all TDMD triggers due to their stringency, they provided a tractable set of promising candidates for initial validation.

### Validation of novel TDMD trigger RNAs

For eight of the enriched miRNAs in our AGO-CLASH experiments, we selected the top three highest ranked candidate triggers for further validation (Fig. 1C, Supplemental Table S1). In one case, miR-450b-5p, only one candidate trigger (*Rdx*) remained after applying all criteria. Given the well-established role of *Cyrano* as a trigger for TDMD of miR-7a-5p (Kleaveland et al. 2018), additional candidates for this miRNA were not tested. As an initial screen, CRISPR interference (CRISPRi) was used to knockdown putative triggers in contact-inhibited immortalized MEFs (Gilbert et al. 2014). Knockdown of *Cyrano* resulted in the expected increase in miR-7a-5p levels without affecting expression of the miR-7 passenger strand (miR-7a-3p or miR-7a*), confirming the ability of this system to detect validated trigger activity (Supplemental Fig. 2A,B). We next knocked down each candidate trigger using two distinct single guide RNAs (sgRNAs) and assessed the levels of the trigger, the target miRNA, and its respective passenger strand using qRT-PCR (Fig. 1D and Supplemental Fig. 2C). In cases where potential triggers were not reliably detected in MEFs (*Syt4*, *Lcp2*, *Ildr2*, *Frem1*, *Gm15477*, *Adam22*, and *Ccr2*), corresponding measurements of the miRNAs were not performed, reasoning that these transcripts could not be responsible for TDMD in this cell type.

Potential triggers for which knockdown with both guides resulted in a statistically significant increase in the respective miRNA without an increase in abundance of the passenger strand were selected for further validation. Importantly, this screening approach may yield false negatives, as residual transcript remaining after CRISPRi-mediated knockdown may be sufficient to carry out TDMD in some cases. Additionally, if multiple redundant triggers exist for a single miRNA, knockdown of one may not be sufficient to result in detectable miRNA de-repression. Nevertheless, even with these caveats, we were able to identify six potential trigger RNAs for four miRNAs of interest (Fig. 1D, Supplemental Fig. 2C). This set included the pairs *Plagl1*:miR-322-5p, *Malat1*:miR-503-5p, *Lrrc58*:miR-503-5p, *Lpar4*:miR-335-3p, *Dnal1*:miR-335-3p, and *Rdx*:miR-450b-5p. Knockdown of the top three candidate triggers for the other miRNAs tested (miR-154-3p, miR-495-3p, miR-485-3p, miR-543-3p and miR-376b-3p) did not reproducibly increase the levels of the corresponding miRNAs (Supplemental Fig. 2C).

To further validate potential triggers supported by CRISPRi experiments, CRISPR/Cas9-mediated genome editing was used to generate clonal MEF cell lines harboring homozygous deletions of each endogenous candidate trigger site (Fig. 2A). These deletions were ∼100 to 300 base pairs in length and were located in 3′ UTRs in all cases except for the site in the noncoding RNA *Malat1*. To test whether loss of the putative trigger site abolished TDMD of the respective miRNA, knockout cell lines were infected with control or *Zswim8*-targeting lentiviral CRISPR vectors and the resulting effects on miRNA expression were assessed. As expected, loss of ZSWIM8 in wild type MEFs led to an increase in the levels of all tested miRNAs, without a corresponding increase in the passenger strands (Fig. 2B-E and Supplemental Fig. 3A-D). In contrast, deletion of multiple individual candidate trigger sites impaired TDMD of their respective miRNAs. Specifically, loss of the miR-322-5p binding site in the 3′ UTR of *Plagl1* fully abrogated regulation of this miRNA by ZSWIM8, providing strong evidence that this transcript represents the sole trigger for miR-322-5p in MEFs (Fig. 2B,C). In the case of miR-503-5p, deletion of the putative trigger site in *Lrrc58* partially inhibited TDMD of this miRNA, while removal of the site in *Malat1* had no detectable effect (Fig. 2D,E). Combined deletion of these trigger sites, however, fully abolished TDMD of miR-503-5p, demonstrating that these transcripts act redundantly to induce decay of this miRNA in MEFs, with *Lrrc58* apparently acting as the dominant trigger. To our knowledge, this represents the first known example of a case in which two TDMD triggers can act on a single miRNA. Of note, human miR-503-5p differs from the mouse sequence by one nucleotide (Supplemental Fig. 3E). Interestingly, there is a corresponding nucleotide change in the putative miR-503-5p binding site in the 3′ UTR of human *LRRC58*, suggesting selective pressure to maintain the base pairing architecture of this site across species.

**Figure 2.**
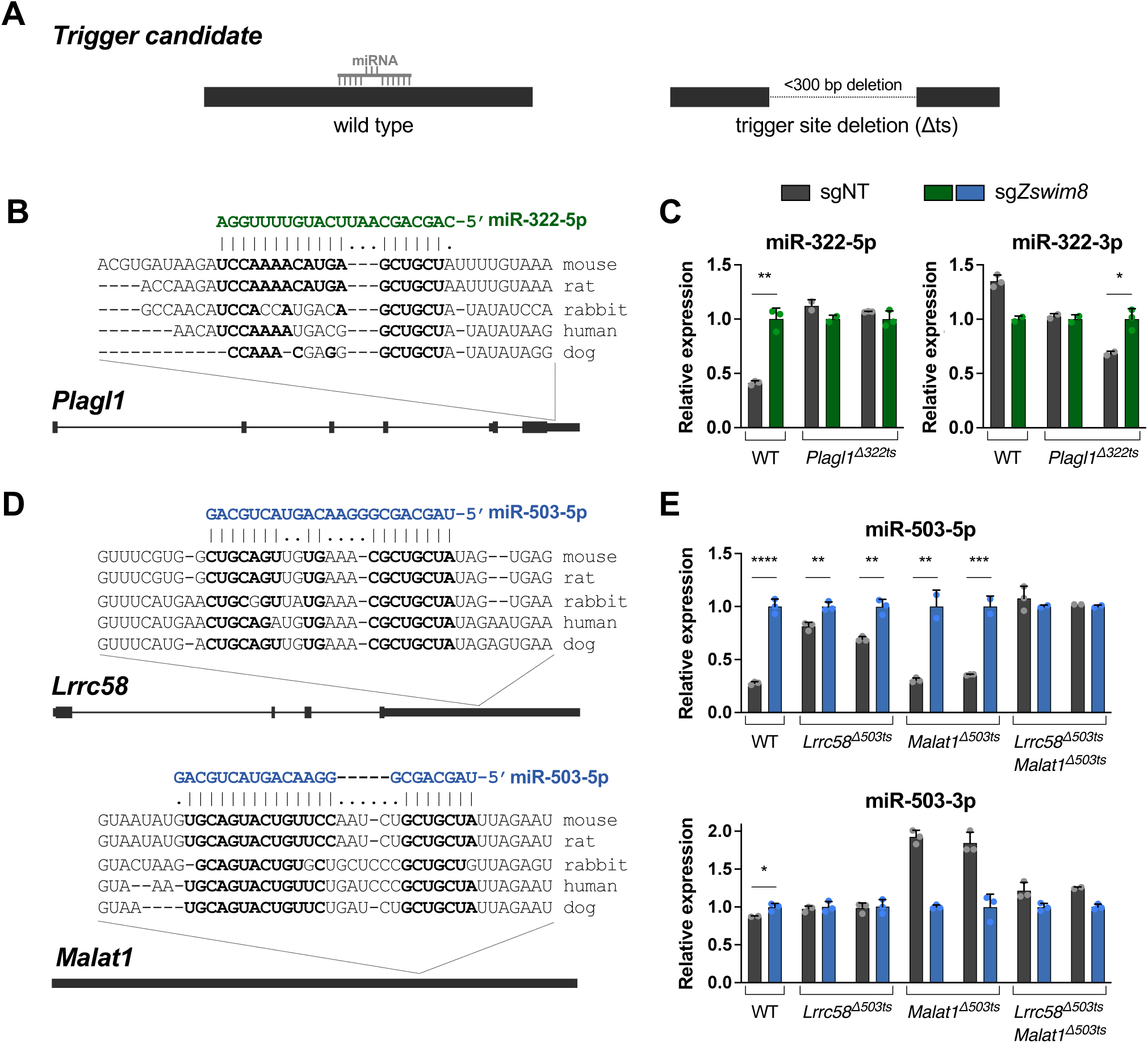
Validation of novel TDMD triggers. (*A*) Schematic of trigger site deletion (Δts) strategy to validate putative TDMD triggers. (*B,D*) Genomic organization of TDMD trigger transcripts with conservation and predicted miRNA base-pairing architecture of the trigger sites. Nucleotides predicted to base pair with the miRNA are shown in bold. (*C,E*) qRT-PCR analysis of indicated miRNAs, relative to miR-16-5p, in WT and Δts MEFs. Parental MEFs or two independent Δts clones for each trigger site were infected with lentivirus expressing Cas9 and a non-targeting CRISPR guide (sgNT) or *Zswim8-*targeting guide (sg*Zswim8*). Values were normalized to expression level in sg*Zswim8* for each condition. n=3 technical replicates per clone with individual data points plotted (mean ± SD shown). *P* values were calculated by one-tailed student’s t-test. \**P*<0.05; \*\**P*<0.01; \*\*\**P*<0.001; \*\*\*\**P*<0.0001.

We also deleted putative trigger sites in *Lpar4* and *Dnal1* for miR-335-3p (alone and in combination), and in *Rdx* for miR-450b-5p, but these deletions did not impair TDMD of these miRNAs (Supplemental Fig. 3A-D). Therefore, these sites either do not function as TDMD triggers or they act redundantly with other sites to induce decay of these miRNAs. Altogether, these experiments identify *Plagl1*, *Lrrc58*, and *Malat1* as functional TDMD triggers in MEFs.

### Mice lacking the TDMD trigger sites in *Plagl1* and *Lrrc58* exhibit embryonic growth restriction

Germline deletion of Z*swim8* in mice results in perinatal lethality, heart and lung defects, and reduced body size at E18.5 (Jones et al. 2023; Shi et al. 2023). Deletion of miR-322 and miR-503 in *Zswim8^—/—^* mice rescued embryonic growth (Jones et al. 2023), strongly suggesting that this aspect of the ZSWIM8-deficiency phenotype is attributable to aberrant upregulation of these miRNAs due to inactivation of TDMD. Interestingly, while the newly-defined miR-503-5p trigger *Lrrc58* has not previously been linked to regulation of embryonic growth in mammals, the miR-322-5p trigger *Plagl1* has been implicated as a regulator of body size in mice (Varrault et al. 2006). *Plagl1* (also known as *Zac1*) is a maternally-imprinted gene that encodes a transcription factor that promotes growth by transactivating *Igf2* expression. A previously generated *Plagl1* knockout allele that eliminates expression of both the open reading frame and the 3′ UTR, where the miR-322-5p trigger site is located, results in embryonic growth restriction in mice, mirroring a key attribute of *Zswim8^—/—^* mice (Varrault et al. 2006). These observations suggested that *Plagl1* may regulate body size both through the activity of the encoded protein and, potentially, the miR-322-5p TDMD trigger site in the mRNA.

To test this possibility, we generated mice with small deletions that remove the trigger sites from the 3′ UTRs of *Plagl1* (referred to as *Plagl1^Δ322ts^*) and *Lrrc58* (*Lrrc58^Δ503ts^*) (Fig. 3A). Because *Plagl1* is a maternally-imprinted gene with exclusive expression of the paternal allele (Spengler et al. 1997; Varrault et al. 2006), functionally heterozygous mice cannot be generated. We therefore denote the *Plagl1* genotype solely based upon the identity of the inherited paternal allele (*Plagl1^+^* or *Plagl1^Δ322ts^*). *Plagl1* and *Lrrc58* are widely expressed in mouse tissues, albeit at highly variable levels (Supplemental Fig. 4A), consistent with their potential to function as triggers for TDMD of miR-322-5p and miR-503-5p, which are broadly regulated by ZSWIM8 (Supplemental Fig. 1A) (Jones et al. 2023; Shi et al. 2023). Notably, alternative splicing in the final exon of *Plagl1* produces transcript variants that contain or lack the miR-322-5p trigger site (Supplemental Fig. 4B).

**Figure 3.**
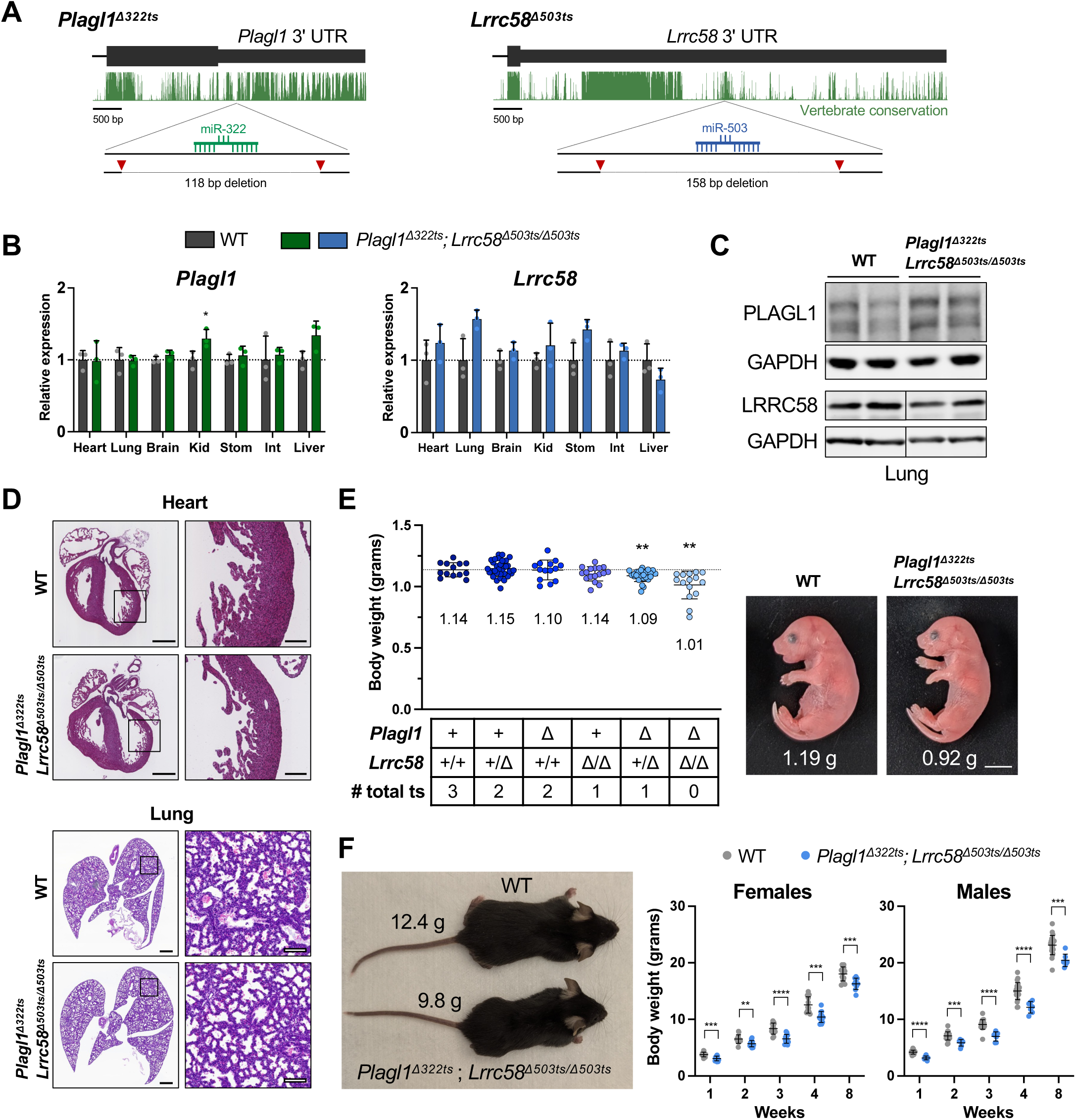
Embryonic growth restriction in *Plagl1^Δ322ts^*; *Lrrc58^Δ503ts/Δ503ts^* mice. (*A*) Schematic of genome-editing strategy to generate *Plagl1^Δ322ts^* and *Lrrc58^Δ503ts^* mice. UCSC genome browser PhastCons 60 vertebrate conservation track shown (mm10). Red triangles depict approximate locations of sgRNAs for CRISPR-mediated editing. (*B*) qRT-PCR analysis of *Plagl1* and *Lrrc58* expression in E18.5 mouse tissues normalized to the geometric mean of two housekeeping genes (*Psmd4* and *Oaz1*). Expression was normalized to mean expression in WT in each tissue. n=3 mice per genotype, with each mouse represented by an individual data point (mean ± SD shown). *P* values were calculated by one-tailed student’s t-test comparing *Plagl1^Δ322ts^; Lrrc58^Δ503ts/Δ503ts^* to WT for each tissue. Kid, kidney; Stom, stomach; Int, small intestine. (*C*) Western blot analysis of PLAGL1 and LRRC58 in E18.5 lung tissues of mice of the indicated genotypes. Irrelevant lanes were removed from blots where indicated with vertical lines. (*D*) Representative hematoxylin and eosin (H&E) stained sections of E18.5 hearts and lungs from mice of the indicated genotypes. n=4 mice were analyzed per genotype. Scale bars are 500 µm (left), 100 µm (right). (*E*) Body weights (left) of E18.5 embryos of the indicated genotypes with each data point representing an individual mouse (mean ±SD shown). Mean weight is denoted on the graph below each cohort. Dotted line is the mean weight of the WT cohort. n=14-32 mice per genotype. *P* values were calculated by one-tailed student’s t-test comparing each genotype to WT. Image (right) of littermate WT and *Plagl1^Δ322ts^; Lrrc58^Δ503ts/Δ503ts^* E18.5 embryos with weights indicated. Scale bar is 0.5 cm. ts, trigger site. (*F*) Image (left) of littermate 4-week-old male WT and *Plagl1^Δ322ts^; Lrrc58^Δ503ts/Δ503ts^* mice with body weights indicated. Graph (right) of body weights of WT and *Plagl1^Δ322ts^; Lrrc58^Δ503ts/Δ503ts^* mice at the indicated timepoints (mean ±SD shown). n=7-22 mice for each genotype at each timepoint. *P* values were calculated by one-tailed student’s t-test. \**P*<0.05; \*\**P*<0.01; \*\*\**P*<0.001; \*\*\*\**P*<0.0001.

In most tissues, the *Plagl1* trigger site deletion did not affect the steady-state abundance of the individual alternatively-spliced *Plagl1* isoforms (Supplemental Fig. 4B) or the overall level of *Plagl1* transcripts (Fig. 3B), although a small increase (<1.5 fold) was observed in selected tissues. A modest but reproducible increase in PLAGL1 protein in lung and heart, but not in MEFs from *Plagl1^Δ322ts^* mice, was also detectable (Fig. 3C and Supplemental Fig. 4C). These findings suggested that the miR-322-5p binding site might confer canonical miRNA-mediated silencing in addition to functioning as a TDMD trigger site in some contexts. The *Lrrc58* trigger site deletion did not significantly affect expression of the mRNA or protein produced from the mutant locus in any tested tissue (Fig. 3B,C and Supplemental Fig. 4C).

Since miR-322-5p and miR-503-5p share the same core 6 nucleotide seed sequence and are therefore expected to share highly overlapping targets, we examined the phenotypes of both single (*Plagl1^Δ322ts^* or *Lrrc58^Δ503ts/Δ503ts^*) and double (*Plagl1^Δ322ts^*; *Lrrc58^Δ503ts/Δ503ts^*) trigger site knockout mice. Mice carrying all combinations of these alleles were viable and were born at the expected Mendelian ratios (Supplemental Fig. 4D). Accordingly, and in contrast to *Zswim8^—/—^* mice, no overt abnormalities of heart or lungs were apparent at E18.5 (Fig. 3D). However, the body size of E18.5 embryos was significantly attenuated in mice carrying trigger site deletions. Specifically, *Plagl1^Δ322ts^*; *Lrrc58^Δ503ts/Δ503ts^*double knockout mice and, to a lesser extent, *Plagl1^Δ322ts^*; *Lrrc58*^+*/Δ503ts*^ mice, exhibited a statistically-significant reduction in embryonic growth (Fig. 3E and Supplemental Fig. 4E). Mice lacking both trigger sites were, on average, ∼11% smaller than wild type controls (Fig. 3E). In comparison, *Zswim8^—/—^* mice were ∼22% smaller than control animals at this developmental time point (Jones et al. 2023; Shi et al. 2023), indicating that stabilization of miR-322-5p and miR-503-5p can account for about half of the growth defect characteristic of ZSWIM8-deficient mice. It is possible that the more severe impairment of embryonic growth in *Zswim8^—/—^* mice is a secondary consequence of the broader developmental abnormalities present in these animals. Alternatively, this effect might be attributable to the upregulation of other TDMD-regulated miRNAs that also function as growth suppressors. Interestingly, while the reduced body size of *Plagl1^Δ322ts^*; *Lrrc58^Δ503ts/Δ503ts^*double knockout mice was maintained into early adulthood (Fig. 3F), the relative postnatal growth rate of these mice was indistinguishable from wild type mice (Supplemental Fig. 4F). This observation suggests that the growth restriction of *Plagl1^Δ322ts^*; *Lrrc58^Δ503ts/Δ503ts^*mice occurs earlier in development, after which mutant animals grow at a normal rate. These results provide strong evidence that the TDMD trigger sites in the 3′ UTRs of *Plagl1* and *Lrrc58* cooperate to control embryonic growth in mice.

### The trigger sites in *Plagl1* and *Lrrc58* mediate TDMD of miR-322-5p and miR-503-5p *in vivo*

We next examined levels of miR-322-5p and miR-503-5p, and their corresponding passenger strands, in tissues from *Plagl1* and *Lrrc58* trigger site knockout mice. Northern blotting of RNA from heart and lung confirmed that loss of the *Plagl1* trigger site specifically de-repressed miR-322-5p, but not its passenger strand (miR-322-3p), while deletion of the *Lrrc58* trigger site resulted in a specific accumulation of miR-503-5p, but not its passenger (miR-503-3p) (Fig. 4A). Analysis of miRNA levels in livers from all combinations of *Plagl1^Δ322ts^* and *Lrrc58^Δ503ts^* genotypes further confirmed the specificity of *Plagl1* and *Lrrc58* trigger sites for their respective miRNAs and documented that the miR-322/503 passenger strands and pri-miRNA were insensitive to trigger site deletions in this tissue (Supplemental Fig. 5A). Similar results were observed in primary MEFs from *Plagl1^Δ322ts^*; *Lrrc58^Δ503ts/Δ503ts^* double knockout mice (Supplemental Fig. 5B). Expanding these analyses to a broad panel of tissues (heart, lung, brain, kidney, stomach, small intestine, and liver) demonstrated that, in all contexts, both miR-322-5p and miR-503-5p were upregulated in *Plagl1^Δ322ts^*; *Lrrc58^Δ503ts/Δ503ts^*mice to a magnitude equivalent to that observed upon knockout of *Zswim8* (Fig. 4B-C). Again, passenger strand levels were unaffected in all analyzed tissues.

**Figure 4.**
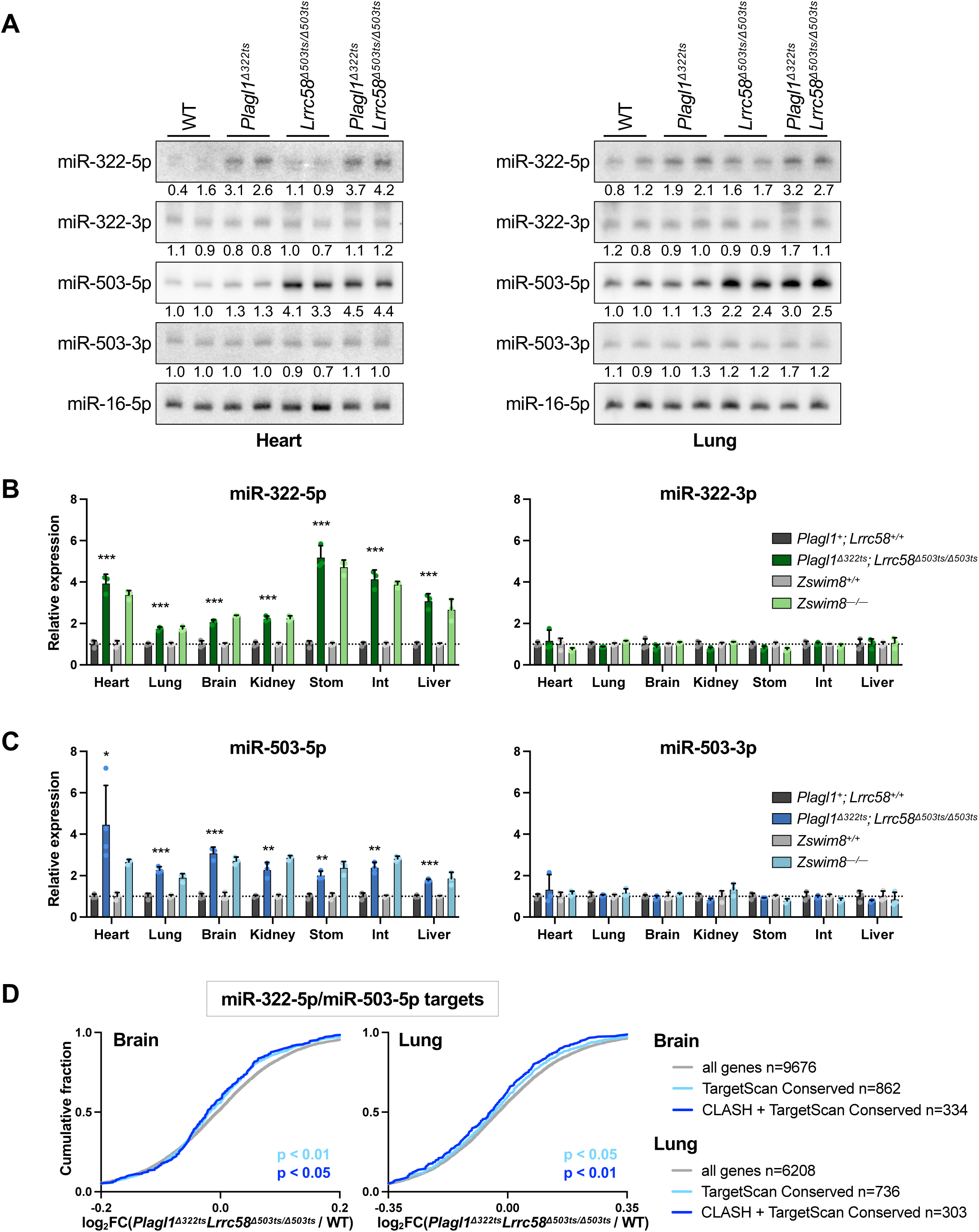
Loss of the *Plagl1* and *Lrrc58* trigger sites abrogates TDMD of miR-322-5p and miR-503-5p in vivo. (*A*) Northern blot analysis of miRNA expression in E18.5 heart (left) and lung (right) in mice of the indicated genotypes. Quantification relative to miR-16-5p, normalized to mean expression in WT, is shown below each lane. (*B,C*) qRT-PCR analysis of miRNAs (left) and passenger strands (right) normalized to miR-16-5p, in mouse tissues of the indicated genotypes at E18.5. Expression in trigger site knockout or *Zswim8* knockout was normalized to mean expression in *Plagl1^+^; Lrrc58^+/+^* or *Zswim8^+/+^* in each tissue, respectively. n=3 mice per genotype, with each mouse represented by an individual data point (mean ± SD shown). *P* values were calculated by one-tailed student’s t-test comparing *Plagl1^Δ322ts^; Lrrc58^Δ503ts/Δ503ts^* to WT for each tissue. Stom, stomach; Int, small intestine. \**P*<0.05; \*\**P*<0.01; \*\*\**P*<0.001. (*D*) Cumulative distribution function (CDF) plot showing the fold-change in expression of the following sets of mRNAs in the brain (left) and lung (right) at E18.5, comparing *Plagl1^Δ322ts^; Lrrc58^Δ503ts/Δ503ts^* to WT: (i) all genes with counts per million (CPM)>5 (gray), (ii) the set of conserved targets of miR-322-5p or miR-503-5p as predicted by TargetScan (light blue) (McGeary et al. 2019), and (iii) the set of conserved TargetScan predicted targets that were also detected as chimeras with miR-322-5p or miR-503-5p in AGO-CLASH experiments (dark blue). *P* value calculated by one-sided Wilcoxon rank sum test.

To determine whether stabilization of miR-322-5p and miR-503-5p in trigger site knockout mice enhances repression of targets of these miRNAs, RNA sequencing was performed on tissues from E18.5 wild type and *Plagl1^Δ322ts^*; *Lrrc58^Δ503ts/Δ503ts^* mice. Consistent with the magnitude of target repression reported previously in *Zswim8^—/—^* tissues (Shi et al. 2023), we observed a modest but significant enhancement of repression of miR-322-5p and miR-503-5p targets in brain and lungs from *Plagl1^Δ322ts^*; *Lrrc58^Δ503ts/Δ503ts^* animals (Fig. 4D). These data therefore establish that *Plagl1* and *Lrrc58* are functional triggers for TDMD of miR-322-5p and miR-503-5p in mouse tissues, and likely represent the sole triggers for these miRNAs in the tissues examined at this developmental time-point (E18.5).

### Growth restriction in *Plagl1* and *Lrrc58* trigger site knockout mice is miR-322/503-dependent

To confirm that trigger site deletions in *Plagl1* and *Lrrc58* restrict embryonic growth in a manner dependent upon miR-322-5p and miR-503-5p, we crossed *Plagl1^Δ322ts^*; *Lrrc58^Δ503ts/Δ503ts^* mice to our previously reported miR-322/503^—/—^ mouse line (Jones et al. 2023). Because of the large number of alleles involved, we specifically compared *Plagl1*^+^; *Lrrc58^Δ503ts/Δ503ts^* mice (whose body size is equivalent to wild type; see Fig. 3E) to *Plagl1^Δ322ts^*; *Lrrc58^Δ503ts/Δ503ts^* mice, with or without an intact miR-322/503 locus present. In keeping with our findings described above, complete loss of the trigger sites resulted in reduced body size of male and female E18.5 embryos in a miR-322/503 wild type background (Fig. 5A-C). As expected, deletion of miR-322/503 resulted in larger embryos, as we previously reported (Jones et al. 2023). Importantly, however, in miR-322/503-deficient mice, complete trigger site deletion did not reduce embryonic growth. Thus, embryonic growth restriction in *Plagl1^Δ322ts^*; *Lrrc58^Δ503ts/Δ503ts^* mice requires miR-322 and miR-503, providing strong evidence that this phenotype is attributable to loss of TDMD of these miRNAs in these animals.

**Figure 5.**
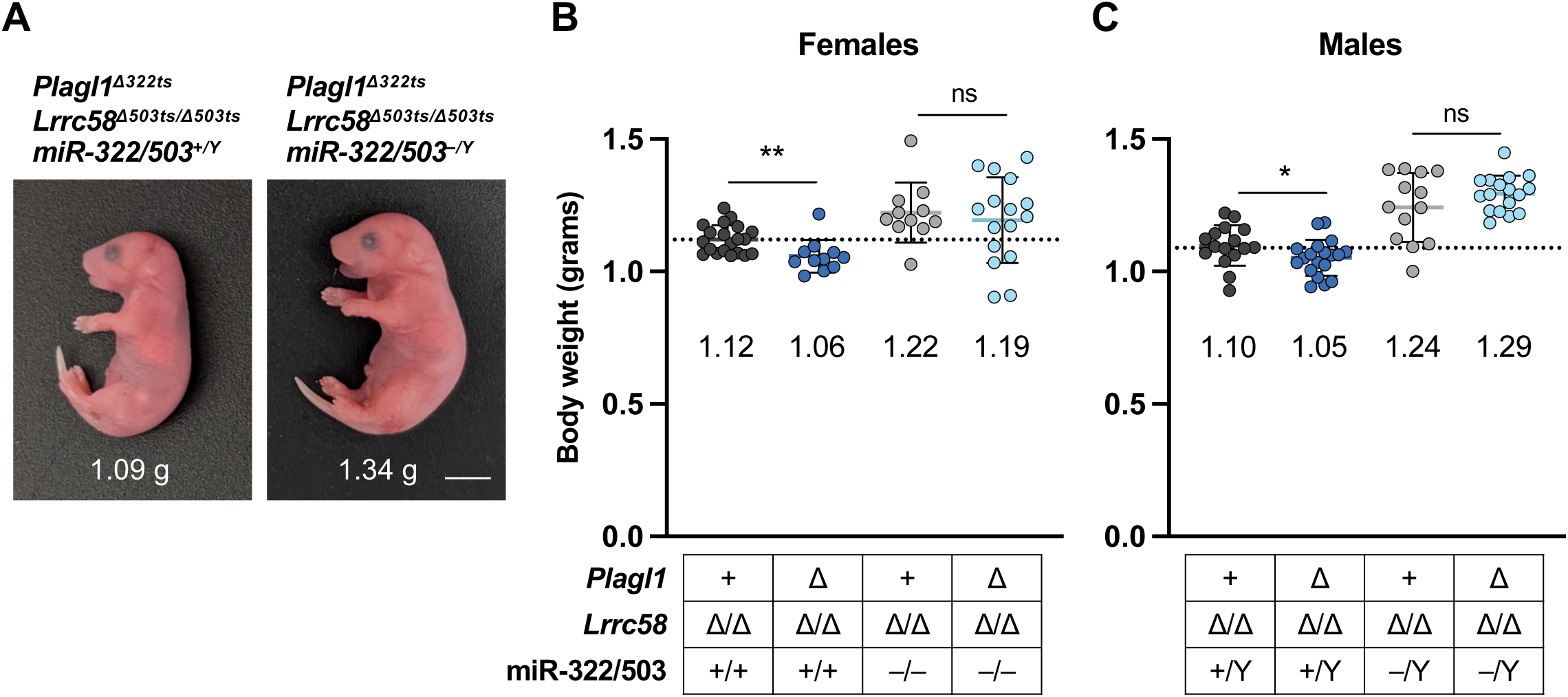
miR-322/503-dependent growth restriction in *Plagl1^Δ322ts^; Lrrc58^Δ503ts/Δ503ts^* mice. (*A*) Images of E18.5 male littermate mice of the indicated genotypes with body weights shown. Scale bar is 1 cm. (*B,C*) Graphs of body weights of female (B) and male (C) E18.5 mice of the indicated genotypes with individual weights plotted (mean ±SD shown). Mean weight is denoted below each cohort. Dotted line is the mean weight of the *Plagl1^+^; Lrrc58^Δ503ts/Δ503ts^* cohort. n=11-20 mice per genotype. *P* values were calculated by one-tailed student’s t-test. \**P*<0.05; \*\**P*<0.01; ns, not significant.

## DISCUSSION

Target-directed microRNA degradation broadly sculpts miRNA expression in metazoans and, accordingly, is essential for normal development in diverse species. Nevertheless, identification of the trigger RNAs that induce decay of many TDMD-regulated miRNAs has proven to be a challenging problem. For example, miR-322-5p and miR-503-5p were among the earliest miRNAs found to exhibit accelerated decay kinetics (Rissland et al. 2011). While ZSWIM8 was later shown to be responsible for the lability of these miRNAs (Shi et al. 2020), the presumptive trigger RNAs that initiate their degradation have remained unknown. Here, using an enhanced AGO-CLASH method (Manakov et al. 2022) focused on the miRNAs that are most strongly regulated by ZSWIM8 in mouse tissues, we identified *Plagl1* and *Lrrc58* as the long-sought TDMD triggers for miR-322-5p and miR-503-5p. Moreover, we demonstrated that loss of these trigger sites in mice results in miR-322/503-dependent embryonic growth restriction, partially recapitulating a central feature of the ZSWIM8-deficiency phenotype. Together, these findings demonstrate that control of miRNA expression by TDMD is essential for normal development in mammals and reveal a noncoding function for the *Plagl1* and *Lrrc58* transcripts as regulators of mammalian body size.

As an initial approach to nominate candidate TDMD trigger sites among the many miRNA-binding sites represented in our AGO-CLASH dataset, we applied a highly-stringent filtering approach that required a perfect seed match and extensive base-pairing between the miRNA 3′ end and the trigger. These criteria were modeled based on features displayed by all previously validated TDMD triggers (Buhagiar and Kleaveland 2024) and enabled identification of the miR-322-5p and miR-503-5p triggers. Nevertheless, this strategy was not sufficient to distinguish *bona fide* triggers from the many possible candidates for other TDMD-regulated miRNAs that were enriched in our AGO-CLASH experiments. There are multiple possible reasons for this limitation that could be addressed in future studies. First, it is likely that the stringent criteria that we applied excluded functional TDMD triggers. Our understanding of the base-pairing architectures that induce TDMD remains incomplete and may vary between different miRNAs. For example, mutagenesis of HSUR1, the herpesvirus-encoded trigger for TDMD of miR-27a, demonstrated that extensive complementarity with the miRNA 3′ end is required for activity of this miRNA:trigger pair (Cazalla et al. 2010; Sheu-Gruttadauria et al. 2019). However, there is at least one example of a miRNA family regulated by the ZSWIM8 homolog EBAX-1 in *C. elegans*, miR-35, whose degradation is specified only by the seed sequence (Donnelly et al. 2022). In this case, mutations in the 3′ end of the miRNA do not impair accelerated miRNA turnover.

While the presumptive trigger that induces degradation of the miR-35 family remains to be identified, these data imply that some TDMD interactions may not require extensive 3′ base-pairing of the miRNA. Such triggers, should they exist in mammals, would likely be represented within our AGO-CLASH dataset but would not be selected as candidate triggers using the bioinformatics approach applied here. Future experiments designed to systematically evaluate the base-pairing requirements for TDMD of individual miRNAs would aid future efforts to rationally nominate candidate trigger sites that engage in empirically-determined base-pairing architectures with their cognate miRNAs.

Redundancy could also potentially confound the identification of new TDMD trigger sites. Indeed, in this study, we identified a case of two triggers acting on the same miRNA, which is, to our knowledge, the first documented example of this principle. While *Lrrc58* is the dominant trigger for TDMD of miR-503-5p, we observed that, at least in MEFs, there is a measurable contribution of a site in the noncoding RNA *Malat1* to TDMD of this miRNA. *Malat1* is a predominantly nuclear transcript, although recent studies have shown it can localize to the cytoplasm in select cell types (Xiao et al. 2024). Regardless, given the extremely high abundance of *Malat1*, localization of even a small fraction of the transcript to the cytoplasm might be sufficient for a productive TDMD interaction in this compartment. In addition, miRNA-loaded AGO proteins can localize to the nucleus and it is possible that TDMD is not limited to cytoplasmic AGO pools (Ohrt et al. 2008; Jeffries et al. 2011; Gagnon et al. 2014; Sarshad et al. 2018; Sala et al. 2023; Johnson et al. 2024). The trigger site that we identified in *Malat1* has previously been shown to bind miR-15-5p and miR-16-5p, which share a core seed sequence with miR-503-5p, in mouse CD8^+^ T cells (Wheeler et al. 2023). Notably, the 3′ sequences of miR-15-5p/16-5p are distinct from miR-503-5p, likely explaining why *Malat1* does not induce TDMD of these miRNAs. Nevertheless, these data provide independent confirmation that this site is accessible to miRNA-mediated AGO binding. This example illustrates the potential for multiple triggers to regulate an individual miRNA, which might necessitate dual knockdown or knockout approaches to uncover the TDMD activity of some yet-to-be validated triggers.

Given our finding that loss of the trigger sites in both *Plagl1* and *Lrrc58* are required to manifest a significant defect in embryonic growth, we can conclude that miR-322-5p and miR-503-5p both function as negative regulators of body size and together contribute to the small body size of ZSWIM8-deficient mice. This is not unexpected in light of the shared core seed sequence and co-regulation of these clustered miRNAs. Together, stabilization of these miRNAs can account for approximately half of the growth defect characteristic of *Zswim8^—/—^* mice (growth defect of ∼11% in *Plagl1^Δ322ts^*; *Lrrc58^Δ503ts/Δ503ts^* mice *vs.* ∼22% in *Zswim8^—/—^* mice) (Jones et al. 2023; Shi et al. 2023). The stronger growth restriction observed in *Zswim8^—/—^*mice could be the result of other developmental abnormalities present in these animals. For example, in humans, congenital heart defects are associated with low birth weight (Aliasi et al. 2023). It also remains possible that stabilization of other TDMD-regulated miRNAs with growth suppressing activity contributes to this phenotype.

Why does accumulation of miR-322-5p and miR-503-5p lead to embryonic growth restriction? These mammalian-specific miRNAs have been reported to regulate diverse pathways, including insulin-like growth factor (IGF) signaling and the cell cycle, both of which play important roles in regulating mammalian body size (Conlon and Raff 1999; Linsley et al. 2007; Rissland et al. 2011; Yang and Xu 2011; Llobet-Navas et al. 2014). For example, *Igf1r*, whose loss results in severe growth restriction in mice (Liu et al. 1993), has been shown to be a target of miR-322-5p and miR-503-5p in mouse mammary epithelial cells (Llobet-Navas et al. 2014). Moreover, as members of the miR-16 family, miR-322-5p and miR-503-5p have the potential to target multiple proteins required for cell cycle progression (Linsley et al. 2007; Bandi et al. 2009; Jiang et al. 2009). Of particular relevance, loss of *Ccnd1*, which encodes cyclin D1 and is a validated target of miR-503-5p and other miR-16 family members, results in small body size in mice (Fantl et al. 1995; Sicinski et al. 1995). Thus, silencing of *Igfr1* and *Ccnd1* offer possible mechanisms linking miR-322-5p/503-5p upregulation to growth restriction. Importantly, despite their smaller body size at birth, the relative postnatal growth rate of *Plagl1^Δ322ts^*; *Lrrc58^Δ503ts/Δ503ts^*mice is indistinguishable from wild type animals. This finding indicates that overexpression of miR-322-5p/503-5p in trigger site knockout mice impacts embryonic growth at an earlier developmental stage, after which mutant animals grow normally. Consistent with this observation, *Plagl1* expression drops dramatically after birth, suggesting that TDMD of miR-322-5p might be less robust in postnatal tissues (Lui et al. 2008). Further study of trigger site-mutant animals throughout embryogenesis is warranted in order to pinpoint the critical developmental window during which accelerated degradation of miR-322-5p/503-5p impacts growth and to identify the specific targets that mediate this effect.

Previous work revealed an example in which the presence of a TDMD trigger site in an mRNA enabled the congruent regulation of a single biological process through dual coding and noncoding functions of the transcript. Specifically, *BCL2L11* promotes apoptosis by encoding the pro-apoptotic protein BIM, while also functioning as a TDMD trigger that removes the apoptosis-suppressing miRNAs miR-221/222 (Zhang et al. 2010; Sionov et al. 2015; Li et al. 2021). Our discovery that the *Plagl1* transcript contains a trigger site for TDMD of miR-322-5p provides another example of this concept. A previously generated knockout allele of *Plagl1*, which eliminates expression of both the protein coding sequence as well as the 3′ UTR where the miR-322-5p trigger site is located, results in fetal growth restriction, altered bone formation, and incompletely penetrant lethality shortly after birth (Varrault et al. 2006). Although the molecular mechanisms underlying these phenotypes are not fully understood, PLAGL1 protein was shown to directly transactivate *Igf2* expression in mice and humans, loss of which likely contributed to growth restriction in knockout animals (Varrault et al. 2006; Iglesias-Platas et al. 2014). Our data reveal a dual role for *Plagl1* in regulating body size, via its canonical protein-coding function as well as through the ability of the transcript to directly eliminate expression of the growth suppressing miRNA miR-322-5p. These dual functions may have contributed to the severity of the *Plagl1* knockout mouse phenotype. Interestingly, alternative splicing in the *Plagl1* 3′ UTR produces two isoforms, one of which lacks the miR-322-5p TDMD trigger site. This configuration provides a mechanism to de-couple production of the PLAGL1 protein from expression of the TDMD trigger. Although we detected broad expression of both *Plagl1* isoforms in bulk mouse tissues at E18.5, it is possible that production of the protein along with high levels of miR-322-5p is advantageous in some settings, which could be achieved by selective expression of the TDMD-deficient *Plagl1* isoform. These observations set the stage for further exploration of the regulation and role of *Plagl1*/*Lrrc58*-mediated degradation of miR-322-5p/miR-503-5p in mammalian body size control.

## MATERIALS AND METHODS

## AGO-CLASH

### Sample preparation

Mouse brain and lung were harvested at E18.5, snap-frozen in liquid nitrogen, and stored at –80 °C. Each sample used for AGO-CLASH consisted of lungs from two mice or a single mouse brain. For UV cross-linking, frozen tissues were ground using a cold mortar and pestle equilibrated in liquid nitrogen, resuspended in 6 mL of 1X Phosphate-Buffered Saline (PBS), and UV cross-linked in a 10 cm tissue culture dish at 243 nm, 400 mJ/cm^2^. Crosslinked tissues were harvested by centrifugation at 400×g for 2 minutes, snap-frozen in liquid nitrogen, and stored at –80 °C until further use.

### AGO immunoprecipitation, chimeric ligation, and sequencing library preparation

For AGO2 immunoprecipitation, 5 μg of AGO2 antibody (eIF2C2 4F9; Santa Cruz) was immobilized on 200 μl of sheep anti-mouse Dynabeads (Invitrogen). Crosslinked lung samples were lysed in 500 μl of iCLIP lysis buffer [50 mM Tris-HCl (pH 7.4), 100 mM NaCl, 1% NP-40, 0.1% SDS, 0.5% sodium deoxycholate, 5.5 μl of 200× Protease Inhibitor Cocktail III (EMD Millipore), 11 μl of Murine RNase inhibitor (New England Biolabs)] and crosslinked brain samples were lysed in 1 ml of iCLIP lysis buffer. Samples were incubated on ice for 5 minutes, followed by sonication using a Bioruptor (Diagenode) on ‘low’ setting at 4 °C for 5 minutes (30 seconds on / 30 seconds off). After lysis, 5 or 10 μl of TurboDNase (Thermo Fisher Scientific) was added to lung or brain samples, respectively, followed by the addition of 10 or 20 μl of 1:300 diluted RNase I (Thermo Fisher Scientific) in PBS, respectively. Lysates were then incubated at 37 °C for 5 minutes with agitation at 1,200 rpm using an Eppendorf ThermoMixer® C. Lysates were cleared by centrifugation at 13,000 × g for 3 minutes at 4 °C, then added to anti-AGO2-immobilized beads. After overnight incubation at 4 °C, bead washes, T4 PNK Minus reaction, RNA chimeric ligation, FastAP treatment, PNK treatment, and ligation of 3′-RNA linker (roJH031 in this study) were conducted as described previously (Manakov et al. 2022). Following proteinase K (New England Biolabs) treatment, RNA isolation was performed using Zymo RNA Clean & Concentrator-5 columns. To enrich specific microRNAs, probe-capture was conducted in brain tissues using oJH1023 (for mmu-miR-322-5p, 503-5p, 154-3p, 543-3p, 450b-5p) and in brain and lung tissues using oJH1024 (for mmu-miR-335-3p, 495-3p, 376b-3p, 7a-5p, 485-3p). Isolated chimeric RNAs from unenriched or enriched samples were subjected to cDNA synthesis using oJH1016/1017 followed by 5′ linker (oJH1018) ligation overnight and then cleanup using Dynabeads MyOne Silane (Thermo Fisher Scientific).

Sequencing libraries were prepared using Q5 PCR master mix (New England Biolabs) with Illumina NextSeq primers and 16-17 PCR cycles. PCR products were separated on a 2% agarose gel and fragments between 190-350 bp in size were extracted from the gel and sequenced on an Illumina NextSeq 2K. At least 50 million reads were obtained for each sample. All oligonucleotide sequences are provided in Supplemental Table S2.

### Sequencing data analysis

UMI barcodes were extracted from the reads using UMI-tools (v1.1.2) (Smith et al. 2017). Adapter sequences were trimmed with Cutadapt (v3.1) (Martin 2011). Reads corresponding to snoRNAs were removed by aligning trimmed reads to mouse snoRNAs using snoRNA Atlas (Jorjani et al. 2016) and Bowtie (v1.3.1) (Langmead et al. 2009). Remaining reads were then mapped to mature mouse miRNA sequences from miRBase v22 (Kozomara et al. 2019) using Bowtie. Reads were filtered based on strand orientation and mismatch rate using chim-eCLIP (https://github.com/YeoLab/chim-eCLIP), which was also used to identify candidate chimeric reads. Chimeric reads that aligned to repetitive elements from the mouse genome (Repbase v27.07) using STAR (v2.7.1a) (Dobin et al. 2013) were excluded from downstream analyses. The remaining reads were aligned to the mouse reference genome (GRCm38) using Bowtie, and duplicate reads were removed using UMI-tools based on UMI barcodes. Enriched peak clusters were called using CLIPper (https://github.com/YeoLab/clipper), with peaks extended by 10 nt at the 5′ end. Annotations were based on GENCODE M23. Conservation scores were calculated using phyloP-60way (Pollard et al. 2010), and the minimum free energy (MFE) of base-pairing between miRNAs and target sequences was calculated using RNAhybrid (Rehmsmeier et al. 2004).

#### Cell culture

HEK293T cells were obtained from ATCC and immortalized MEFs were generated previously (Kopp et al. 2019). Cells were cultured in DMEM (high glucose, pyruvate) supplemented with 10% FBS (Sigma) and either 1X penicillin-streptomycin antibiotic (Thermo Fisher Scientific) or 1X Antibiotic-Antimycotic (Invitrogen). Cell lines were confirmed to be free of mycoplasma contamination.

#### Primary MEF generation

Primary MEFs were isolated from E14.5 mouse embryos as described (Tan and Lei 2019). Briefly, E14.5 embryos were dissected and the head, liver and heart were removed. The remaining tissue was finely minced in cold 0.25% Trypsin (Gibco), pipetted up and down ten times with a 5 mL serological pipette, and then incubated for 10 minutes at 37 °C. Cells were then strained through a 40 μm cell strainer, spun at 500 g 4 °C for 5 minutes, resuspended in 5 mL fresh DMEM with 10% (v/v) fetal bovine serum (Sigma) and 1×Antibiotic-Antimycotic (Invitrogen), and plated in a 60 mm tissue culture dish. Primary MEFs were grown for less than 5 passages.

#### Lentiviral CRISPR and CRISPRi

Immortalized MEFs stably expressing dCas9-KRAB were generated by infecting cells with lentiviral construct pHR-SFFV-dCas9-BFP-KRAB (Addgene #46911) (Gilbert et al. 2014). Single cells positive for BFP were sorted into 96 well plates using a Melody cell sorter (BD Biosciences) and clonal cell lines were established. sgRNAs targeting candidate trigger RNAs, as well as non-target (NT) sgRNAs (sequences provided in Supplemental Table S2), were cloned into pUC6-sgRNA EF1Alpha-puro-T2A-BFP (Addgene #60955) (Gilbert et al. 2014).

Guide sequences were taken from the mCRISPRiv2 library (Horlbeck et al. 2016) or designed using CRISPick (Doench et al. 2016; Sanson et al. 2018). For pooled *Zswim8* knockout experiments, previously reported lentiCRISPRv2-hygro vectors expressing non-targeting or *Zswim8*-targeting guides were used (Han et al. 2020).

To generate lentivirus, 0.5-1x10^6^ HEK293T cells were seeded per well in a 6-well plate. 24 hours after seeding, cells were transfected with 500 ng lentiviral vector, 300 ng psPAX2 (Addgene #12260), and 200 ng pMD2.G (Addgene#12259) using 3 µL FugeneHD (Promega) transfection reagent. At 24 hours post transfection, the medium was changed and 48 hours after transfection, viral supernatant was collected and passed through a 0.45 µm filter. Recipient cells were transduced with viral supernatant in the presence of 8 µg/mL polybrene (Millipore).

Beginning 48 hours after transduction, cells were selected in medium containing 2 µg/mL puromycin (Invivogen) or 200-400 µg/mL hygromycin B (Invitrogen) for 5 days. MEFs were contact-inhibited for 1-3 days prior to harvesting.

#### Generation of trigger site knockout cell lines

MEF trigger site knockout clones were generated in one of two ways: (1) Guide sequences were cloned into px458 (Addgene #48138) (Ran et al. 2013) expressing GFP and Cas9, and immortalized MEFs were transfected with plasmids using FuGeneHD (Promega). 24 hours after transfection, GFP positive cells were sorted into 96 well plates using a Melody cell sorter (BD Biosciences) and clonal cell lines were established. (2) sgRNAs were synthesized (Integrated DNA Technologies), resuspended at a final concentration of 25 µM, and incubated with 25 µM Alt-R S.p. Cas9 Nuclease V3 (Integrated DNA Technologies) in PBS for 15 minutes at room temperature. Electroporation Enhancer (IDT) was added to a final concentration of 12 µM along with 0.6 µg pMax GFP. The mixture was electroporated into immortalized MEFs using the Amaxa P3 Primary Cell kit (Lonza) with program EH-100 on a 4D Nucleofector X unit (Lonza). 72-96 hours after nucleofection, GFP positive cells were sorted into 96 well plates and clonal cell lines were established. Cell lines were screened for deletions by PCR and amplicon sequencing. Amplicon sequencing was analyzed using CRISPResso2 (Clement et al. 2019). sgRNA sequences are provided in Supplemental Table S2.

#### RNA isolation and qRT-PCR

Total RNA was isolated using the miRNeasy Mini Kit (Qiagen) with DNase I digestion according to the manufacturer’s instructions. For lysing cells, Qiazol (Qiagen) was added directly to tissue culture plates after rinsing with PBS. For E18.5 tissues, organs were homogenized in Qiazol (Qiagen) using a Precellys Evolution homogenizer (Bertin Technologies). cDNA was synthesized according to the manufacturer’s instructions using Mir-X miRNA First-Strand Synthesis kit (Takara) for miRNAs and U6, or PrimeScript RT (Takara) for all other transcripts. qRT-PCR was performed with SYBR Green Master Mix (Applied Biosystems) on a QuantStudio 3 (Applied Biosystems). miR-16 or U6 snRNA was used to normalize qRT-PCR measurements of miRNA expression. *Actb* was used to normalize mRNA measurements in cell culture experiments, while the geometric mean of *Psmd4* and *Oaz1* was used to normalize mRNA expression in tissues (Rehfeld et al. 2018). Primer sequences are provided in Supplemental Table S2.

#### Generation and husbandry of genetically engineered mice

All mouse experiments were approved by the University of Texas Southwestern Medical Center Animal Care and Use Committee and performed in accordance with National Institutes of Health guidelines (animal protocol 2017-102001). Mice were group housed under conventional conditions in a 12 hr day/night cycle with normal chow diet (Harlan Teklad TD2916) and water provided ad libitum. *Zswim8^—/—^* (Jax strain #039272) and miR-332/503*^—/—^*mice (Jax strain #039271) were previously described (Jones et al. 2023).

*Plagl1^Δ322ts^; Lrrc58^Δ503ts/Δ503ts^* mice were generated in the University of Texas Southwestern Transgenic Core by microinjection of Cas9 in complex with sgRNAs (Integrated DNA Technologies) into the pronucleus of fertilized C57BL/6J eggs. Sequences of sgRNAs are provided in Supplemental Table S2. Founders carrying the desired alleles were maintained on a pure C57BL/6J background and backcrossed continuously. For timed matings, the morning of detection of vaginal plug was defined as E0.5.

#### Western Blotting

Cells were lysed in RIPA buffer (150 mM NaCl,1% NP-40, 0.5% sodium deoxycholate, 0.1% SDS, 25 mM Tris-HCl pH 8.0), with 2×EDTA free protease inhibitor cocktail (Roche) and 1 mM phenylmethylsulfonyl fluoride (Sigma). Mouse tissues were lysed in the same buffer using a Bioruptor Plus (Diagenode). Lysates were cleared by centrifugation. Proteins were separated by SDS-PAGE and transferred to nitrocellulose membranes (0.2 µm, Cytiva). Blots were blocked in TBS with 0.1% Tween-20 (TBST) containing 5% non-fat milk and incubated with primary antibodies in TBST containing 3% BSA overnight at 4 °C. After washing in TBST, the blots were incubated with secondary antibodies in TBST with 3% BSA for one hour at room temperature followed by further washes. Images were acquired on an Odyssey fluorescent imaging system (LI-COR Biosystems). Primary antibodies used were: anti-HA (1:1000 Cell Signaling C29F4), anti-PLAGL1 (1:500 Santa Cruz #166944), anti-LRRC58 (1:100 ThermoFisher # PA5-63453), and anti-GAPDH (1:2000 Santa Cruz #32233). Secondary antibodies used were anti-Mouse and anti-Rabbit secondary IR Dye 800CW (LI-COR).

#### Histology

Tissues were fixed in 10% formalin, embedded in paraffin, and sectioned to show max surface area of the lung or 4-chambered view of the heart. Hematoxylin and eosin (H&E) staining was performed on paraffin sections. Slides were scanned using a Leica Aperio slide scanner and images generated in Aperio ImageScope (Leica Biosystems).

#### Northern blotting

RNA isolation was performed as described above. For heart samples, 2-3 hearts of the same genotype were pooled for each lane. 20 μg of RNA was separated on 15% TBE-Urea polyacrylamide gels. RNAs were transferred to BrightStar-Plus nylon membranes (Invitrogen) followed by UV-crosslinking at 120 mJ/cm^2^. Membranes were pre-hybridized with ULTRAhyb-Oligo hybridization buffer (Invitrogen) followed by hybridization with ^32^P end-labeled oligonucleotide probes. A DNA oligonucleotide probe was used for miR-16-5p, while miRCURY LNA miRNA Detection probes (Qiagen) were used for miR-322-5p, miR-322-3p, miR-503-5p, and miR-503-3p. Probe sequences are provided in Supplementary Table S2. Densitometry was performed using Quantity One 1-D Analysis software (Bio-Rad), and miR-16-5p was used as a normalization control.

#### RNA sequencing

RNA-seq libraries were constructed using the QuantSeq 3′ mRNA-Seq V2 Library Prep Kit FWD with UDI 12 nt Set A1 and B1 (Lexogen, 192.24) using 500 ng input RNA. Libraries were sequenced on a NextSeq 2K (Illumina) with 65 bp single-end reads. Reads were mapped to the reference mouse genome (GRCm39) using STAR (v2.7.11b) (Dobin et al. 2013). Gene-level read counts were obtained using featureCounts (v1.6.0) (Liao et al. 2014) with Gencode M31 annotation (Frankish et al. 2023). Differential expression analysis was performed using the edgeR package (v4.2.1) (Chen et al. 2025).

#### Conservation Alignments

For base level conservation analyses (Figure 2 and Supplemental Figure 3), sequences were obtained from the following genome assemblies: *Mus musculus* GRCm38/mm10, *Rattus norvegicus* RGSC 6.0/rn6, *Oryctolagus cuniculus* Broad/oryCun2, *Homo sapiens* GRCh38/hg38, *Canis lupus familiaris* UU_Cfam_GSD_1.0/canFam4. Conservation tracks were obtained from the UCSC genome browser (PhastCons 60 vertebrate conservation track) (Yang 1995; Felsenstein and Churchill 1996; Siepel et al. 2005; Pollard et al. 2010).

#### Statistics

To determine statistical significance, one-tailed studentʹs t test and chi-square test were calculated using GraphPad Prism (version 10.5.0) and Wilcoxon rank sum test with continuity correction was calculated using R. Values are reported as mean ± SD in all figures.

## ACKNOWLEDGEMENTS

We thank Feng Zhang, Jonathan Weissman, and Didier Trono for plasmids. We thank Robert Hammer and the UT Southwestern Transgenic core, John Shelton and the UT Southwestern Histo Pathology Core, and Vanessa Schmid and the McDermott Center Next Generation Sequencing Core. We thank Jeanetta Marshburn-Wynn for assistance with mouse husbandry, Frederick Rehfeld for technical assistance, and Kathryn O’Donnell and members of the Mendell laboratory for helpful suggestions on the manuscript. This work was supported by CPRIT (RP220309 to J.T.M.), the Welch Foundation (I-1961-20240404 to J.T.M.), NIH (R01CA282036 to J.T.M.), a National Research Foundation of Korea (NRF) grant funded by the Korean Ministry of Science (MSIT) (RS-2024-00335939 and RS-2025-00556082 to J.H.), and the Brain Pool Program funded by the Korean Ministry of Science (MIST) and Information and Communication Technology (ICT) through the National Research Foundation of Korea (NRF) (RS-2024-00399517 to M.K.). J.T.M. is an Investigator of the Howard Hughes Medical Institute.

## AUTHOR CONTRIBUTIONS

C.A.L., J.H., S.C., M.K., B.M.E., T-C.C., A.A., and J.T.M. designed experiments and interpreted the results. C.A.L., J.H., S.C., M.K., K.S., T-C.C., and A.A. performed the experiments. H.Z. and J.H. performed bioinformatic analyses. C.A.L., J.H., and J.T.M. wrote the manuscript.

## REFERENCES

Alberti C, Cochella L. 2017. A framework for understanding the roles of miRNAs in animal development. Development 144: 2548–2559.

Aliasi M, Mastenbroek M, Papakosta S, van Geloven N, Haak MC. 2023. Birthweight of children with isolated congenital heart disease—A sibling analysis study. Prenatal Diagnosis 43: 639–646.

Ameres SL, Horwich MD, Hung JH, Xu J, Ghildiyal M, Weng Z, Zamore PD. 2010. Target RNA-directed trimming and tailing of small silencing RNAs. Science 328: 1534–1539.

Bail S, Swerdel M, Liu H, Jiao X, Goff LA, Hart RP, Kiledjian M. 2010. Differential regulation of microRNA stability. RNA 16: 1032–1039.

Bandi N, Zbinden S, Gugger M, Arnold M, Kocher V, Hasan L, Kappeler A, Brunner T, Vassella E. 2009. miR-15a and miR-16 are implicated in cell cycle regulation in a Rb-dependent manner and are frequently deleted or down-regulated in non-small cell lung cancer. Cancer Res 69: 5553–5559.

Bartel DP. 2018. Metazoan MicroRNAs. Cell 173: 20–51.

Bitetti A, Mallory AC, Golini E, Carrieri C, Carreño Gutiérrez H, Perlas E, Pérez-Rico YA, Tocchini-Valentini GP, Enright AJ, Norton WHJ et al. 2018. MicroRNA degradation by a conserved target RNA regulates animal behavior. Nat Struct Mol Biol 25: 244–251.

Buhagiar AF, Kleaveland B. 2024. To kill a microRNA: emerging concepts in target-directed microRNA degradation. Nucleic Acids Res 52: 1558–1574.

Cazalla D, Yario T, Steitz JA. 2010. Down-regulation of a host microRNA by a Herpesvirus saimiri noncoding RNA. Science 328: 1563–1566.

Chen Y, Chen L, Lun ATL, Baldoni PL, Smyth GK. 2025. edgeR v4: powerful differential analysis of sequencing data with expanded functionality and improved support for small counts and larger datasets. Nucleic Acids Res 53: gkaf018.

Clement K, Rees H, Canver MC, Gehrke JM, Farouni R, Hsu JY, Cole MA, Liu DR, Joung JK, Bauer DE et al. 2019. CRISPResso2 provides accurate and rapid genome editing sequence analysis. Nat Biotechnol 37: 224–226.

Conlon I, Raff M. 1999. Size Control in Animal Development. Cell 96: 235–244.

DeVeale B, Swindlehurst-Chan J, Blelloch R. 2021. The roles of microRNAs in mouse development. Nat Rev Genet 22: 307–323.

Dobin A, Davis CA, Schlesinger F, Drenkow J, Zaleski C, Jha S, Batut P, Chaisson M, Gingeras TR. 2013. STAR: ultrafast universal RNA-seq aligner. Bioinformatics 29: 15–21.

Doench JG, Fusi N, Sullender M, Hegde M, Vaimberg EW, Donovan KF, Smith I, Tothova Z, Wilen C, Orchard R et al. 2016. Optimized sgRNA design to maximize activity and minimize off-target effects of CRISPR-Cas9. Nat Biotechnol 34: 184–191.

Donnelly BF, Yang B, Grimme AL, Vieux KF, Liu CY, Zhou L, McJunkin K. 2022. The developmentally timed decay of an essential microRNA family is seed-sequence dependent. Cell Rep 40: 111154.

Fantl V, Stamp G, Andrews A, Rosewell I, Dickson C. 1995. Mice lacking cyclin D1 are small and show defects in eye and mammary gland development. Genes Dev 9: 2364–2372.

Felsenstein J, Churchill GA. 1996. A Hidden Markov Model approach to variation among sites in rate of evolution. Mol Biol Evol 13: 93–104.

Frankish A, Carbonell-Sala S, Diekhans M, Jungreis I, Loveland JE, Mudge JM, Sisu C, Wright JC, Arnan C, Barnes I et al. 2023. GENCODE: reference annotation for the human and mouse genomes in 2023. Nucleic Acids Res 51: D942–d949.

Friedman RC, Farh KK, Burge CB, Bartel DP. 2009. Most mammalian mRNAs are conserved targets of microRNAs. Genome Res 19: 92–105.

Gagnon KT, Li L, Chu Y, Janowski BA, Corey DR. 2014. RNAi factors are present and active in human cell nuclei. Cell Rep 6: 211–221.

Gantier MP, McCoy CE, Rusinova I, Saulep D, Wang D, Xu D, Irving AT, Behlke MA, Hertzog PJ, Mackay F et al. 2011. Analysis of microRNA turnover in mammalian cells following Dicer1 ablation. Nucleic Acids Res 39: 5692–5703.

Gatfield D, Le Martelot G, Vejnar CE, Gerlach D, Schaad O, Fleury-Olela F, Ruskeepää AL, Oresic M, Esau CC, Zdobnov EM et al. 2009. Integration of microRNA miR-122 in hepatic circadian gene expression. Genes Dev 23: 1313–1326.

Gebert LFR, MacRae IJ. 2019. Regulation of microRNA function in animals. Nat Rev Mol Cell Biol 20: 21–37.

Ghini F, Rubolino C, Climent M, Simeone I, Marzi MJ, Nicassio F. 2018. Endogenous transcripts control miRNA levels and activity in mammalian cells by target-directed miRNA degradation. Nat Commun 9: 3119.

Gilbert LA, Horlbeck MA, Adamson B, Villalta JE, Chen Y, Whitehead EH, Guimaraes C, Panning B, Ploegh HL, Bassik MC et al. 2014. Genome-Scale CRISPR-Mediated Control of Gene Repression and Activation. Cell 159: 647–661.

Grimson A, Farh KK, Johnston WK, Garrett-Engele P, Lim LP, Bartel DP. 2007. MicroRNA targeting specificity in mammals: determinants beyond seed pairing. Mol Cell 27: 91–105.

Guo Y, Liu J, Elfenbein SJ, Ma Y, Zhong M, Qiu C, Ding Y, Lu J. 2015. Characterization of the mammalian miRNA turnover landscape. Nucleic Acids Res 43: 2326–2341.

Han J, LaVigne CA, Jones BT, Zhang H, Gillett F, Mendell JT. 2020. A ubiquitin ligase mediates target-directed microRNA decay independently of tailing and trimming. Science 370: eabc9546.

Helwak A, Tollervey D. 2014. Mapping the miRNA interactome by cross-linking ligation and sequencing of hybrids (CLASH). Nat Protoc 9: 711–728.

Hiers NM, Li T, Traugot CM, Xie M. 2024. Target-directed microRNA degradation: Mechanisms, significance, and functional implications. Wiley Interdiscip Rev RNA 15: e1832.

Horlbeck MA, Gilbert LA, Villalta JE, Adamson B, Pak RA, Chen Y, Fields AP, Park CY, Corn JE, Kampmann M et al. 2016. Compact and highly active next-generation libraries for CRISPR-mediated gene repression and activation. Elife 5: e19760.

Hwang HW, Wentzel EA, Mendell JT. 2007. A hexanucleotide element directs microRNA nuclear import. Science 315: 97–100.

Iglesias-Platas I, Martin-Trujillo A, Petazzi P, Guillaumet-Adkins A, Esteller M, Monk D. 2014. Altered expression of the imprinted transcription factor PLAGL1 deregulates a network of genes in the human IUGR placenta. Hum Mol Genet 23: 6275–6285.

Jeffries CD, Fried HM, Perkins DO. 2011. Nuclear and cytoplasmic localization of neural stem cell microRNAs. RNA 17: 675–686.

Jiang Q, Feng MG, Mo YY. 2009. Systematic validation of predicted microRNAs for cyclin D1. BMC Cancer 9: 194.

Johnson KC, Kilikevicius A, Hofman C, Hu J, Liu Y, Aguilar S, Graswich J, Han Y, Wang T, Westcott JM et al. 2024. Nuclear localization of Argonaute 2 is affected by cell density and may relieve repression by microRNAs. Nucleic Acids Res 52: 1930–1952.

Jonas S, Izaurralde E. 2015. Towards a molecular understanding of microRNA-mediated gene silencing. Nat Rev Genet 16: 421–433.

Jones BT, Han J, Zhang H, Hammer RE, Evers BM, Rakheja D, Acharya A, Mendell JT. 2023. Target-directed microRNA degradation regulates developmental microRNA expression and embryonic growth in mammals. Genes Dev 37: 661–674.

Jorjani H, Kehr S, Jedlinski DJ, Gumienny R, Hertel J, Stadler PF, Zavolan M, Gruber AR. 2016. An updated human snoRNAome. Nucleic Acids Res 44: 5068–5082.

Kim H, Lee YY, Kim VN. 2025. The biogenesis and regulation of animal microRNAs. Nat Rev Mol Cell Biol 26: 276–296.

Kingston ER, Bartel DP. 2019. Global analyses of the dynamics of mammalian microRNA metabolism. Genome Res 29: 1777–1790.

Kingston ER, Blodgett LW, Bartel DP. 2022. Endogenous transcripts direct microRNA degradation in Drosophila, and this targeted degradation is required for proper embryonic development. Mol Cell 82: 3872–3884.e3879.

Kleaveland B, Shi CY, Stefano J, Bartel DP. 2018. A Network of Noncoding Regulatory RNAs Acts in the Mammalian Brain. Cell 174: 350–362.e317.

Kopp F, Elguindy MM, Yalvac ME, Zhang H, Chen B, Gillett FA, Lee S, Sivakumar S, Yu H, Xie Y et al. 2019. PUMILIO hyperactivity drives premature aging of Norad-deficient mice. eLife 8: e42650.

Kozomara A, Birgaoanu M, Griffiths-Jones S. 2019. miRBase: from microRNA sequences to function. Nucleic Acids Res 47: D155–D162.

Krol J, Busskamp V, Markiewicz I, Stadler MB, Ribi S, Richter J, Duebel J, Bicker S, Fehling HJ, Schübeler D et al. 2010. Characterizing light-regulated retinal microRNAs reveals rapid turnover as a common property of neuronal microRNAs. Cell 141: 618–631.

Langmead B, Trapnell C, Pop M, Salzberg SL. 2009. Ultrafast and memory-efficient alignment of short DNA sequences to the human genome. Genome Biol 10: R25.

Lewis BP, Burge CB, Bartel DP. 2005. Conserved seed pairing, often flanked by adenosines, indicates that thousands of human genes are microRNA targets. Cell 120: 15–20.

Li L, Sheng P, Li T, Fields CJ, Hiers NM, Wang Y, Li J, Guardia CM, Licht JD, Xie M. 2021. Widespread microRNA degradation elements in target mRNAs can assist the encoded proteins. Genes Dev 35: 1595–1609.

Li T, Li L, Hiers NM, Sheng P, Wang Y, Traugot CM, Effinger-Morris JF, Akaphan P, Liu Y, Bian J et al. 2025. Translation suppresses exogenous target RNA-mediated microRNA decay. Nat Commun 16: 5257.

Liao Y, Smyth GK, Shi W. 2014. featureCounts: an efficient general purpose program for assigning sequence reads to genomic features. Bioinformatics 30: 923–930.

Linsley PS, Schelter J, Burchard J, Kibukawa M, Martin MM, Bartz SR, Johnson JM, Cummins JM, Raymond CK, Dai H et al. 2007. Transcripts targeted by the microRNA-16 family cooperatively regulate cell cycle progression. Mol Cell Biol 27: 2240–2252.

Liu JP, Baker J, Perkins AS, Robertson EJ, Efstratiadis A. 1993. Mice carrying null mutations of the genes encoding insulin-like growth factor I (Igf-1) and type 1 IGF receptor (Igf1r). Cell 75: 59–72.

Llobet-Navas D, Rodríguez-Barrueco R, Castro V, Ugalde AP, Sumazin P, Jacob-Sendler D, Demircan B, Castillo-Martín M, Putcha P, Marshall N et al. 2014. The miR-424(322)/503 cluster orchestrates remodeling of the epithelium in the involuting mammary gland. Genes Dev 28: 765–782.

Lui JC, Finkielstain GP, Barnes KM, Baron J. 2008. An imprinted gene network that controls mammalian somatic growth is down-regulated during postnatal growth deceleration in multiple organs. Am J Physiol Regul Integr Comp Physiol 295: R189–196.

Manakov SA, Shishkin AA, Yee BA, Shen KA, Cox DC, Park SS, Foster HM, Chapman KB, Yeo GW, Van Nostrand EL. 2022.Scalable and deep profiling of mRNA targets for individual microRNAs with chimeric eCLIP. bioRxiv doi: 10.1101/2022.1102.1113.480296.

Martin M. 2011. Cutadapt removes adapter sequences from high-throughput sequencing reads. J Comput Biol 24: 1138–1143.

Marzi MJ, Ghini F, Cerruti B, de Pretis S, Bonetti P, Giacomelli C, Gorski MM, Kress T, Pelizzola M, Muller H et al. 2016. Degradation dynamics of microRNAs revealed by a novel pulse-chase approach. Genome Res 26: 554–565.

McGeary SE, Lin KS, Shi CY, Pham TM, Bisaria N, Kelley GM, Bartel DP. 2019. The biochemical basis of microRNA targeting efficacy. Science 366: eaav1741.

Mendell JT, Olson EN. 2012. MicroRNAs in stress signaling and human disease. Cell 148: 1172–1187.

Molina-Pelayo C, Olguin P, Mlodzik M, Glavic A. 2022. The conserved Pelado/ZSWIM8 protein regulates actin dynamics by promoting linear actin filament polymerization. Life Sci Alliance 5: e202201484.

Ohrt T, Mütze J, Staroske W, Weinmann L, Höck J, Crell K, Meister G, Schwille P. 2008. Fluorescence correlation spectroscopy and fluorescence cross-correlation spectroscopy reveal the cytoplasmic origination of loaded nuclear RISC in vivo in human cells. Nucleic Acids Res 36: 6439–6449.

Perez G, Barber GP, Benet-Pages A, Casper J, Clawson H, Diekhans M, Fischer C, Gonzalez JN, Hinrichs AS, Lee CM et al. 2025. The UCSC Genome Browser database: 2025 update. Nucleic Acids Res 53: D1243–D1249.

Pollard KS, Hubisz MJ, Rosenbloom KR, Siepel A. 2010. Detection of nonneutral substitution rates on mammalian phylogenies. Genome Res 20: 110–121.

Ran FA, Hsu PD, Wright J, Agarwala V, Scott DA, Zhang F. 2013. Genome engineering using the CRISPR-Cas9 system. Nat Protoc 8: 2281–2308.

Rehfeld F, Maticzka D, Grosser S, Knauff P, Eravci M, Vida I, Backofen R, Wulczyn FG. 2018. The RNA-binding protein ARPP21 controls dendritic branching by functionally opposing the miRNA it hosts. Nat Commun 9: 1235.

Rehmsmeier M, Steffen P, Hochsmann M, Giegerich R. 2004. Fast and effective prediction of microRNA/target duplexes. RNA 10: 1507–1517.

Reichholf B, Herzog VA, Fasching N, Manzenreither RA, Sowemimo I, Ameres SL. 2019. Time-Resolved Small RNA Sequencing Unravels the Molecular Principles of MicroRNA Homeostasis. Mol Cell 75: 756–768.e757.

Rissland OS, Hong SJ, Bartel DP. 2011. MicroRNA destabilization enables dynamic regulation of the miR-16 family in response to cell-cycle changes. Mol Cell 43: 993–1004.

Sala L, Kumar M, Prajapat M, Chandrasekhar S, Cosby RL, La Rocca G, Macfarlan TS, Awasthi P, Chari R, Kruhlak M et al. 2023. AGO2 silences mobile transposons in the nucleus of quiescent cells. Nat Struct Mol Biol 30: 1985–1995.

Sanson KR, Hanna RE, Hegde M, Donovan KF, Strand C, Sullender ME, Vaimberg EW, Goodale A, Root DE, Piccioni F et al. 2018. Optimized libraries for CRISPR-Cas9 genetic screens with multiple modalities. Nat Commun 9: 5416.

Sarshad AA, Juan AH, Muler AIC, Anastasakis DG, Wang X, Genzor P, Feng X, Tsai PF, Sun HW, Haase AD et al. 2018. Argonaute-miRNA Complexes Silence Target mRNAs in the Nucleus of Mammalian Stem Cells. Mol Cell 71: 1040–1050.e1048.

Shang R, Lee S, Senavirathne G, Lai EC. 2023. microRNAs in action: biogenesis, function and regulation. Nat Rev Genet 24: 816–833.

Sheng P, Li L, Li T, Wang Y, Hiers NM, Mejia JS, Sanchez JS, Zhou L, Xie M. 2023. Screening of Drosophila microRNA-degradation sequences reveals Argonaute1 mRNA’s role in regulating miR-999. Nat Commun 14: 2108.

Sheu-Gruttadauria J, Pawlica P, Klum SM, Wang S, Yario TA, Schirle Oakdale NT, Steitz JA, MacRae IJ. 2019. Structural Basis for Target-Directed MicroRNA Degradation. Mol Cell 75: 1243–1255.e1247.

Shi CY, Elcavage LE, Chivukula RR, Stefano J, Kleaveland B, Bartel DP. 2023. ZSWIM8 destabilizes many murine microRNAs and is required for proper embryonic growth and development. Genome Res 33: 1482–1496.

Shi CY, Kingston ER, Kleaveland B, Lin DH, Stubna MW, Bartel DP. 2020. The ZSWIM8 ubiquitin ligase mediates target-directed microRNA degradation. Science 370: eabc9359.

Sicinski P, Donaher JL, Parker SB, Li T, Fazeli A, Gardner H, Haslam SZ, Bronson RT, Elledge SJ, Weinberg RA. 1995. Cyclin D1 provides a link between development and oncogenesis in the retina and breast. Cell 82: 621–630.

Siepel A, Bejerano G, Pedersen JS, Hinrichs AS, Hou M, Rosenbloom K, Clawson H, Spieth J, Hillier LW, Richards S et al. 2005. Evolutionarily conserved elements in vertebrate, insect, worm, and yeast genomes. Genome Res 15: 1034–1050.

Simeone I, Rubolino C, Noviello Teresa Maria R, Farinello D, Cerulo L, Marzi Matteo J, Nicassio F. 2022. Prediction and pan-cancer analysis of mammalian transcripts involved in target directed miRNA degradation. Nucleic Acids Research 50: 2019–2035.

Sionov RV, Vlahopoulos SA, Granot Z. 2015. Regulation of Bim in Health and Disease. Oncotarget 6: 23058–23134.

Smith T, Heger A, Sudbery I. 2017. UMI-tools: modeling sequencing errors in Unique Molecular Identifiers to improve quantification accuracy. Genome Res 27: 491–499.

Spengler D, Villalba M, Hoffmann A, Pantaloni C, Houssami S, Bockaert J, Journot L. 1997. Regulation of apoptosis and cell cycle arrest by Zac1, a novel zinc finger protein expressed in the pituitary gland and the brain. EMBO J 16: 2814–2825.

Stubna MW, Shukla A, Bartel DP. 2024. Widespread destabilization of Caenorhabditis elegans microRNAs by the E3 ubiquitin ligase EBAX-1. RNA 31: 51–66.

Tan YS, Lei YL. 2019. Generation and Culture of Mouse Embryonic Fibroblasts. Methods Mol Biol 1960: 85–91.

van Rooij E, Sutherland LB, Qi X, Richardson JA, Hill J, Olson EN. 2007. Control of stress-dependent cardiac growth and gene expression by a microRNA. Science 316: 575–579.

Varrault A, Gueydan C, Delalbre A, Bellmann A, Houssami S, Aknin C, Severac D, Chotard L, Kahli M, Le Digarcher A et al. 2006. Zac1 regulates an imprinted gene network critically involved in the control of embryonic growth. Dev Cell 11: 711–722.

Wang G, Lei J, Wang Y, Yu J, He Y, Zhao W, Hu Z, Xu Z, Jin Y, Gu Y et al. 2022. The ZSWIM8 ubiquitin ligase regulates neurodevelopment by guarding the protein quality of intrinsically disordered Dab1. Cerebral Cortex 33: 3866–3881.

Wang Z, Hou Y, Guo X, van der Voet M, Boxem M, Dixon JE, Chisholm AD, Jin Y. 2013. The EBAX-type Cullin-RING E3 ligase and Hsp90 guard the protein quality of the SAX-3/Robo receptor in developing neurons. Neuron 79: 903–916.

Wheeler BD, Gagnon JD, Zhu WS, Muñoz-Sandoval P, Wong SK, Simeonov DS, Li Z, DeBarge R, Spitzer MH, Marson A et al. 2023. The lncRNA Malat1 inhibits miR-15/16 to enhance cytotoxic T cell activation and memory cell formation. Elife 12: RP87900.

Winter J, Jung S, Keller S, Gregory RI, Diederichs S. 2009. Many roads to maturity: microRNA biogenesis pathways and their regulation. Nat Cell Biol 11: 228–234.

Xiao W, Halabi R, Lin CH, Nazim M, Yeom KH, Black DL. 2024. The lncRNA Malat1 is trafficked to the cytoplasm as a localized mRNA encoding a small peptide in neurons. Genes Dev 38: 294–307.

Yang X, Xu T. 2011. Molecular mechanism of size control in development and human diseases. Cell Res 21: 715–729.

Yang Z. 1995. A space-time process model for the evolution of DNA sequences. Genetics 139: 993–1005.

Zhang CZ, Zhang JX, Zhang AL, Shi ZD, Han L, Jia ZF, Yang WD, Wang GX, Jiang T, You YP et al. 2010. MiR-221 and miR-222 target PUMA to induce cell survival in glioblastoma. Mol Cancer 9: 229.

