## Supplemental Figures for "*Plagl1* and *Lrrc58* control mammalian body size by triggering target-directed microRNA degradation of miR-322 and miR-503"

### Supplemental Figure 1

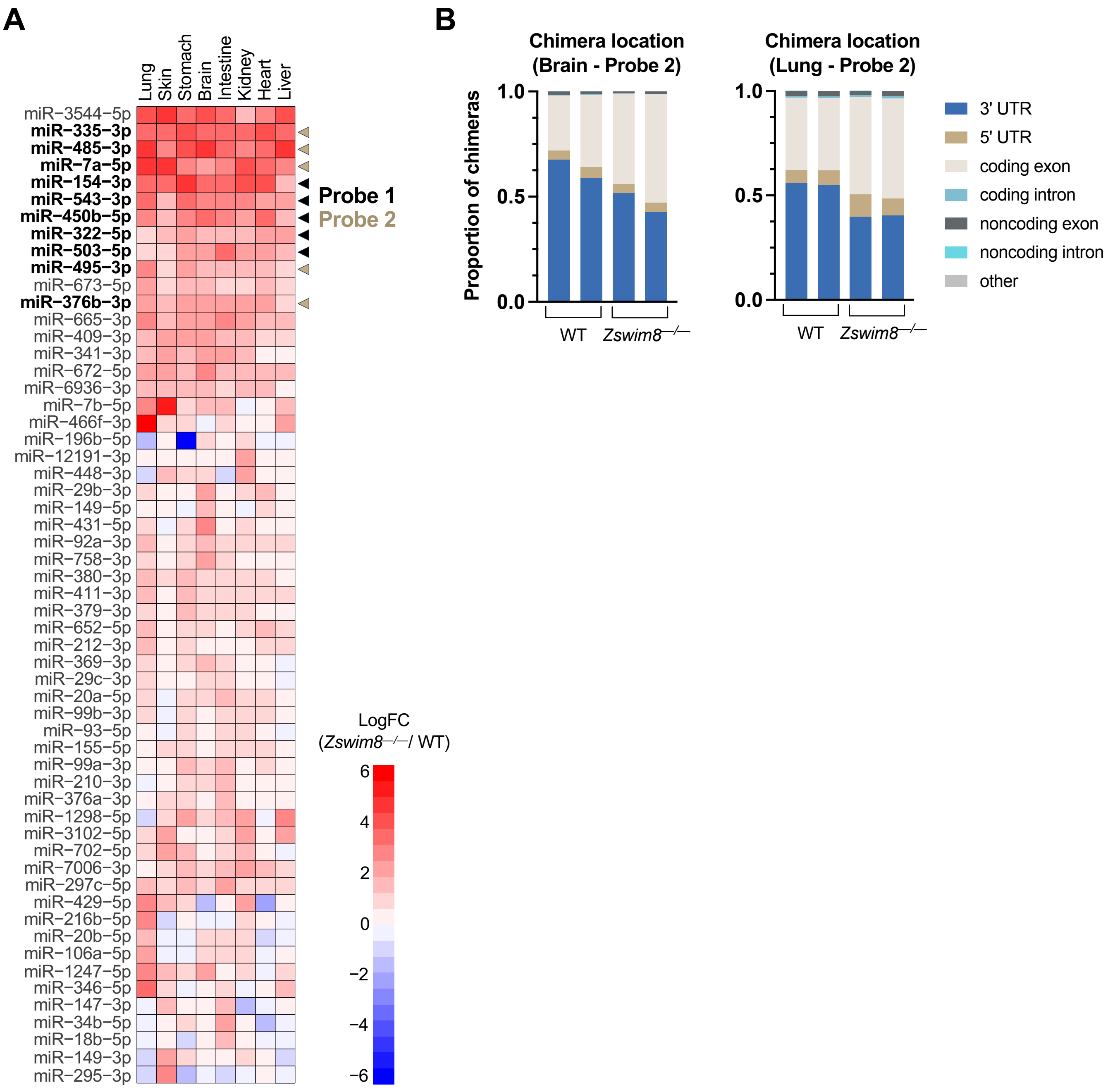


**Supplemental Figure 1. miRNA expression in *Zswim8*$^{-/-}$ mice and analysis of AGO-CLASH data, related to Figure 1.** (*A*) Heat map showing previously reported $\log_2$ fold change (LogFC) of miRNA expression in E18.5 *Zswim8*$^{-/-}$ versus WT tissues (Jones et al. 2023). miRNAs that were enriched with probes for AGO-CLASH experiments are indicated with arrowheads. (*B*) Proportion of chimeras mapped to each location in replicate WT and *Zswim8*$^{-/-}$ brain (left) and lung (right) AGO-CLASH samples enriched with Probe 2. Only chimeras in which the predicted base pairing to the miRNA seed sequence was a 6mer, 7mer-A1, 7mer-m8, or 8mer were included in the analysis.

### Supplemental Figure 2

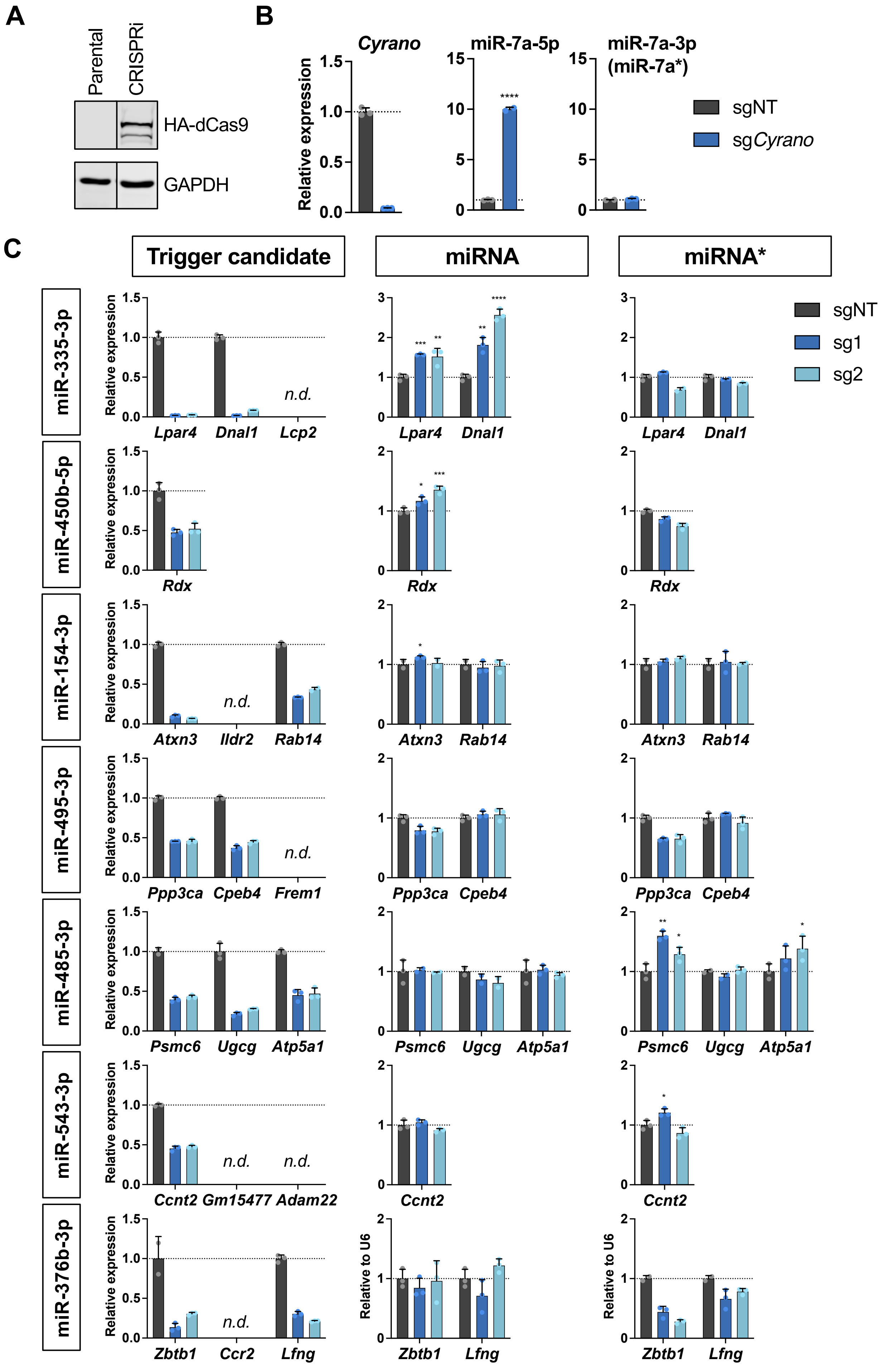

**Supplemental Figure 2. CRISPRi-based screening of candidate TDMD trigger RNAs, related to Figure 1.** (*A*) Western blot of parental MEFs and a clonal MEF cell line expressing dCas9-HA-BFP-KRAB. GAPDH is shown as loading control. Irrelevant lanes were removed from blots where indicated with vertical lines. (*B,C*) dCas9-KRAB-expressing immortalized MEFs were infected with lentivirus encoding a non-targeting guide (sgNT), a guide targeting *Cyrano* (B), or two independent guides targeting each indicated candidate trigger (sg1 or sg2; panel C). Candidate trigger expression was normalized to *Actb* (left), mature miRNA abundance was normalized to miR-16-5p (middle), and passenger strand (miRNA*) levels were normalized to miR-16-5p (right) except for miR-376b-3p and miR-376b-5p, which were normalized to U6. Values were normalized to expression level in sgNT for each transcript. n=3 technical replicates per sgRNA with individual data points plotted (mean ± SD shown). *P* values were calculated by one-tailed student's t-test comparing sg1 or sg2 to sgNT. *$P<0.05$; **$P<0.01$; ***$P<0.001$; ****$P<0.0001$; n.d., not reliably detected.

#### Supplemental Figure 3

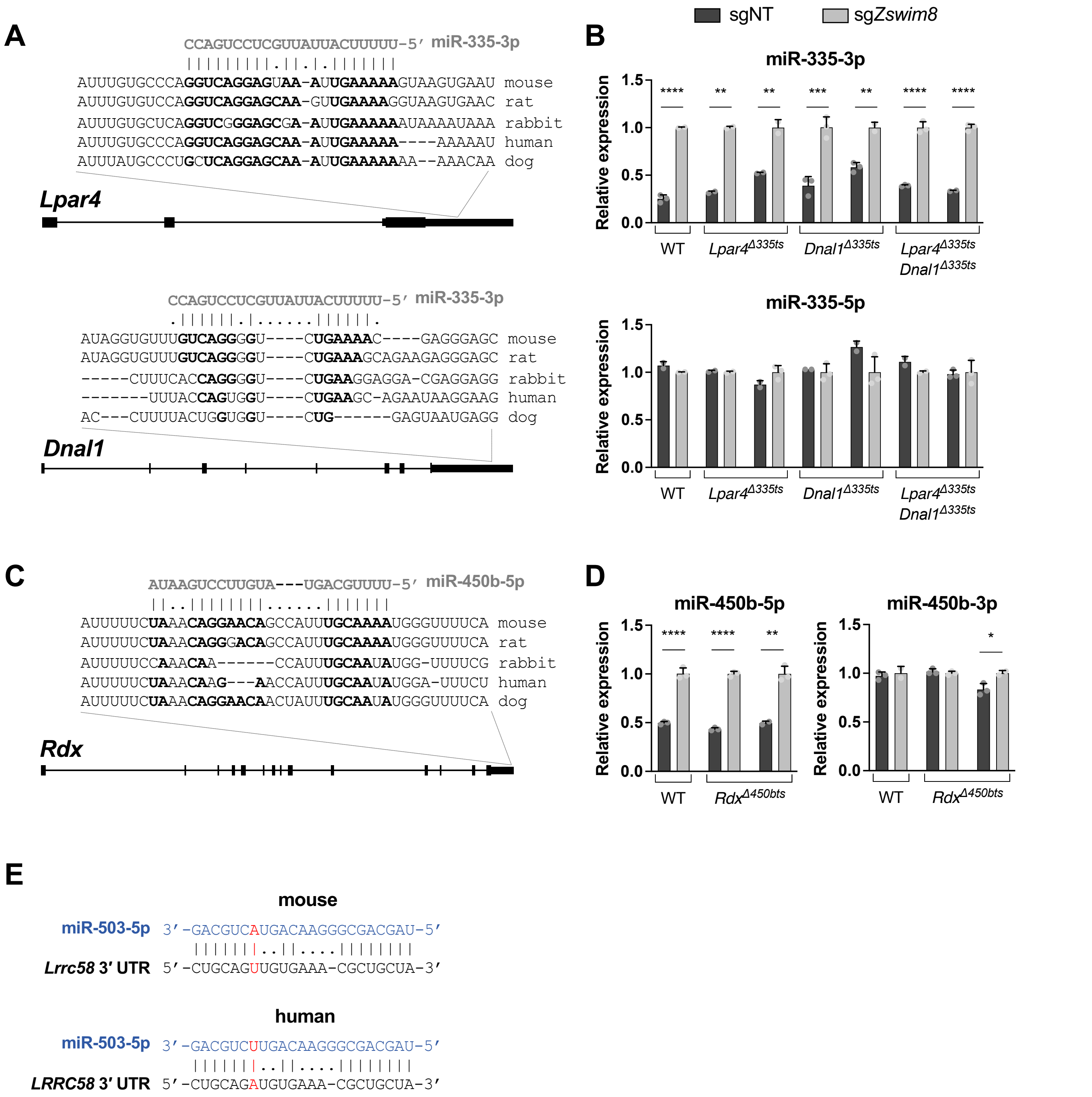


**Supplemental Figure 3. Deletion of additional candidate TDMD trigger sites, related to Figure 2.** (*A,C*) Genomic organization of candidate TDMD trigger transcripts with conservation and predicted miRNA base-pairing architecture of the trigger sites. Nucleotides predicted to base pair with the miRNA are shown in bold. (*B,D*) qRT-PCR analysis of indicated miRNAs, normalized to miR-16-5p, in WT and Δts MEFs. Parental MEFs or two independent Δts clones for each candidate trigger site were infected with lentivirus expressing Cas9 and a non-targeting CRISPR guide (sgNT) or *Zswim8* targeting guide (sg*Zswim8*). Values were normalized to expression level in sg*Zswim8* for each condition. n=3 technical replicates per clone with individual data points plotted (mean ± SD shown). *P* values were calculated by one-tailed student's t-test. *$P<0.05$; **$P<0.01$; ***$P<0.001$; ****$P<0.0001$. (*E*) Predicted base pairing of miR-503-5p to the TDMD trigger site in mouse *Lrrc58* (top) and in human *LRRC58* (bottom). Nucleotides that differ between mouse and human are shown in red.

### Supplemental Figure 4

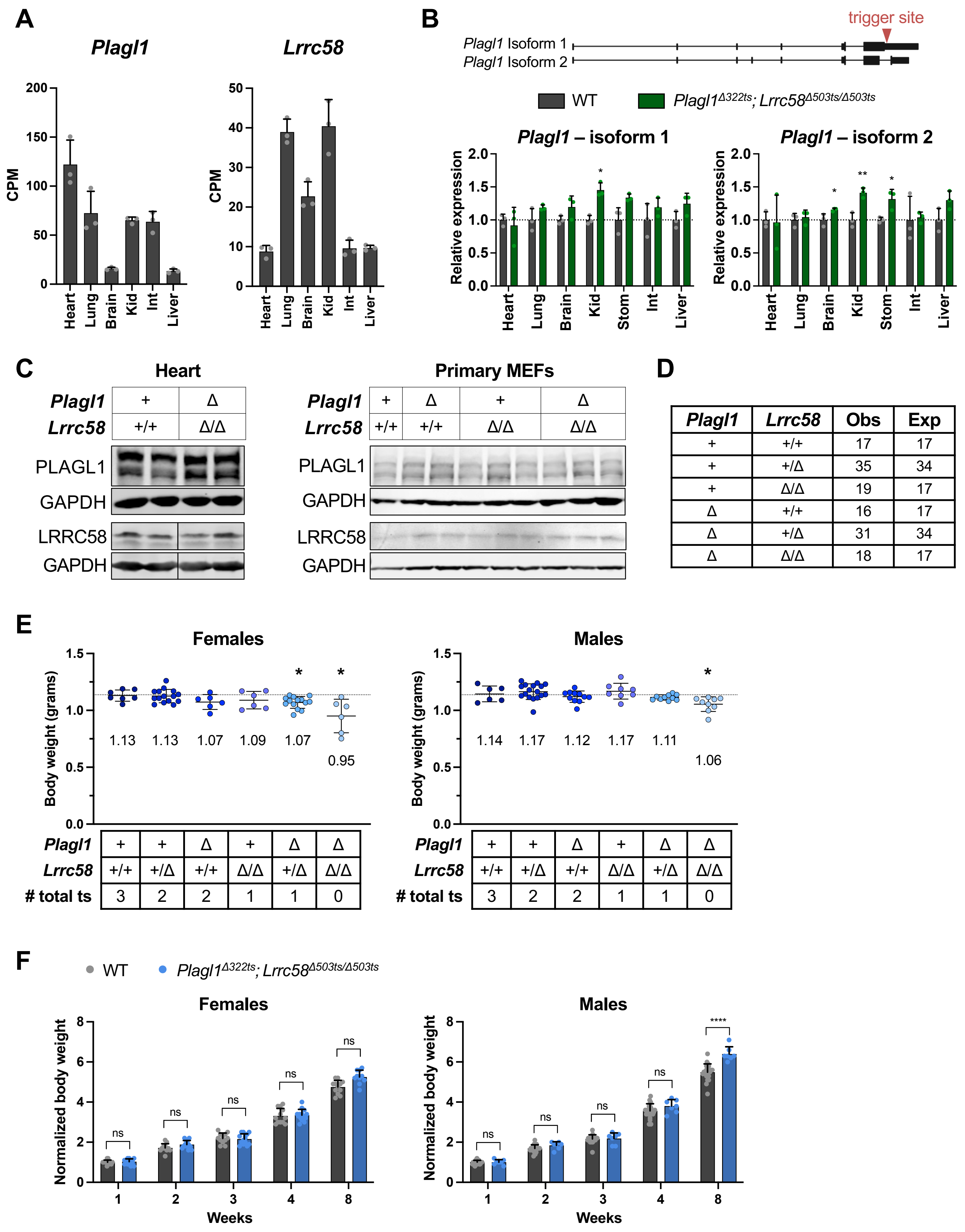

**Supplemental Figure 4.** ***Plagl1$^{\Delta 322ts}$*; *Lrrc58$^{\Delta 503ts/\Delta 503ts}$* mice exhibit reduced body size but normal post-natal growth rate, related to Figure 3.** (*A*) Expression in counts per million (CPM) of *Plagl1* and *Lrrc58* in WT mouse tissues at E18.5, measured by RNA-seq. (*B*) qRT-PCR analysis of alternatively-spliced *Plagl1* isoforms in mouse tissues of the indicated genotypes, normalized to the geometric mean of two housekeeping genes (*Psmd4* and *Oaz1*). Expression was normalized to mean expression in WT in each tissue. Isoform 1 includes the miR-322-5p trigger site in the 3′ UTR, which is spliced out in isoform 2. n=3 mice per genotype, with each mouse represented by an individual data point (mean ± SD shown). *P* values were calculated by one-tailed student's t-test comparing *Plagl1$^{\Delta 322ts}$*; *Lrrc58$^{\Delta 503ts/\Delta 503ts}$* to WT for each tissue. (*C*) Western blot analysis of PLAGL1 and LRRC58 in E18.5 hearts and in primary MEFs from embryos of the indicated genotypes. GAPDH is shown as a loading control. Irrelevant lanes were removed from blots where indicated with vertical lines. (*D*) Table showing numbers of observed (Obs) and expected (Exp) mice of the indicated genotypes resulting from a *Plagl1$^{+/\Delta 322ts}$*; *Lrrc58$^{+/\Delta 503ts}$* intercross. Chi-squared=0.647, df=5; two-tailed *P* value=0.995. (*E*) Body weights of female (left) and male (right) E18.5 embryos of the indicated genotypes with each data point representing an individual mouse (mean ±SD shown). Mean weight is denoted on the graph below each cohort. Dotted line is the mean weight of the WT cohort. n=6-17 mice per genotype. *P* values were calculated by one-tailed student's t-test comparing each genotype to WT. ts, trigger site. (*F*) Graph of body weights of WT and *Plagl1$^{\Delta 322ts}$*; *Lrrc58$^{\Delta 503ts/\Delta 503ts}$* mice at the indicated timepoints normalized to the average weight of each respective genotype at 1-week of age (mean ±SD shown). n=7-22 mice for each genotype at each timepoint. *P* values were calculated by one-tailed student's t-test. **P*<0.05; ***P*<0.01; *****P*<0.0001; ns, not significant.

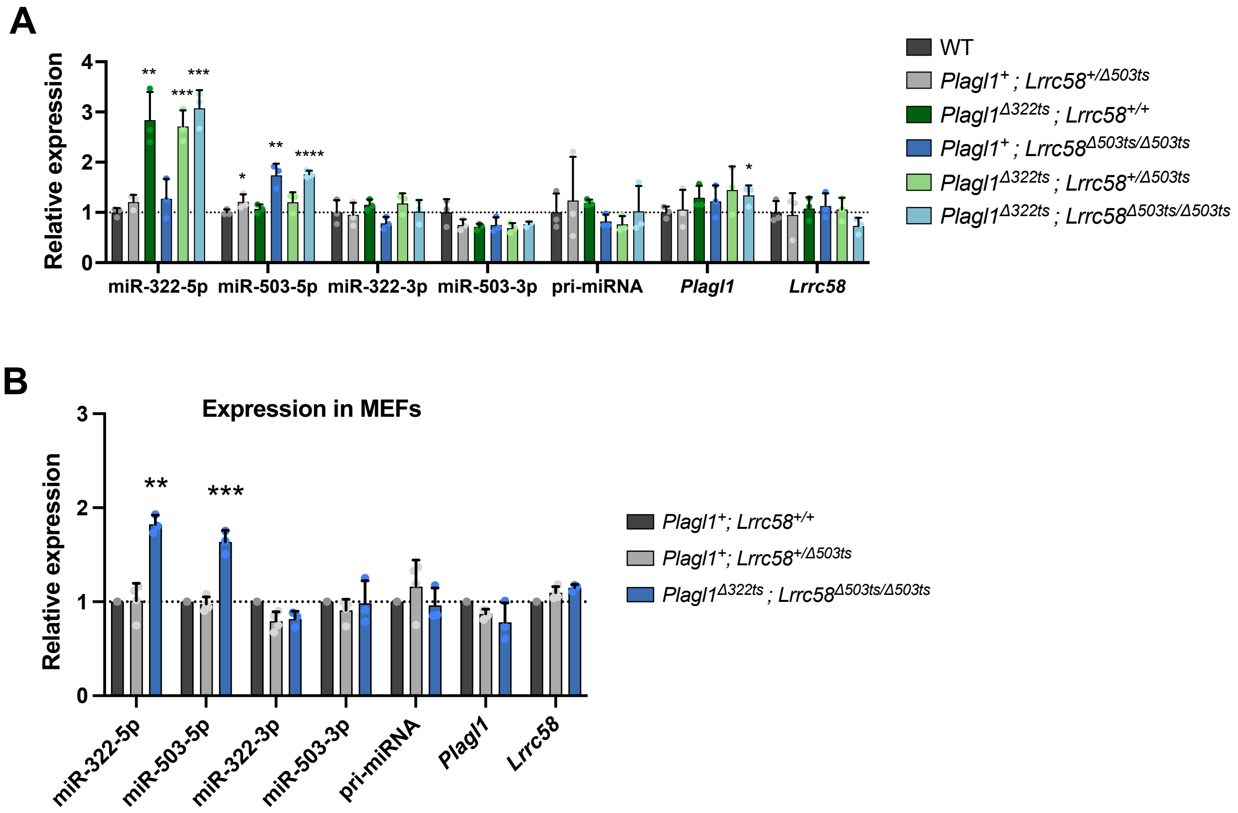


**Supplemental Figure 5. Loss of the *Plagl1* and *Lrrc58* trigger sites abrogates TDMD of miR-322-5p and miR-503-5p *in vivo*, related to Figure 4.** (*A*) qRT-PCR analysis of the indicated transcripts in E18.5 livers. Expression of each transcript was normalized to mean expression in WT. n=3 biological replicates per genotype with individual data points shown (mean ± SD shown). *P* values were calculated by one-tailed student's t-test comparing each genotype to WT. (*B*) qRT-PCR analysis of the indicated transcripts in primary MEFs. Expression of each transcript was normalized to expression in $Plagl1^{+}$; $Lrrc58^{+/+}$. n=1 biological replicate for $Plagl1^{+}$; $Lrrc58^{+/+}$, n=4 biological replicates for $Plagl1^{+}$; $Lrrc58^{+/\Delta 503ts}$, and n=3 biological replicates for $Plagl1^{\Delta 322ts}$; $Lrrc58^{\Delta 503ts/\Delta 503ts}$ with individual data points shown (mean ± SD shown). *P* values were calculated by one-tailed student's t-test comparing $Plagl1^{\Delta 322ts}$; $Lrrc58^{\Delta 503ts/\Delta 503ts}$ to $Plagl1^{+}$; $Lrrc58^{+/\Delta 503ts}$. *$P<0.05$; **$P<0.01$; ***$P<0.001$; ****$P<0.0001$.
